# EGFR inhibition led ROCK activation enhances desmosome assembly and cohesion in cardiomyocytes

**DOI:** 10.1101/2022.04.27.489705

**Authors:** Maria Shoykhet, Orsela Dervishi, Philipp Menauer, Matthias Hiermaier, Colin Osterloh, Ralf J. Ludwig, Jens Waschke, Sunil Yeruva

## Abstract

Arrhythmogenic cardiomyopathy (AC) is a familial heart disease partly caused by impaired desmosome turnover. Thus, stabilization of desmosome integrity may provide potential new treatment options. Desmosomes, apart from cellular cohesion, provide the structural framework of a signaling hub. Here, we investigated the role of the epidermal growth factor receptor (EGFR) in cardiomyocyte cohesion. We inhibited EGFR under physiological and pathophysiological conditions using the murine plakoglobin knockout AC model, in which EGFR was upregulated. EGFR inhibition enhanced cardiomyocyte cohesion. Immunoprecipitation showed an interaction of EGFR and desmoglein 2 (DSG2). Immunostaining and AFM revealed enhanced DSG2 localization and binding at cell borders upon EGFR inhibition. Enhanced area composita length and desmosome assembly were observed upon EGFR inhibition, confirmed by enhanced DSG2 and desmoplakin (DP) recruitment to cell borders. Erlotinib, an EGFR inhibitor, activated ROCK. Erlotinib mediated desmosome assembly and cardiomyocyte cohesion were abolished upon ROCK inhibition. Thus, inhibiting EGFR, thereby stabilizing desmosome integrity, might provide new treatment options for AC.

**Summary:** Shoykhet *et al*. show that EGFR inhibition led ROCK activation enhances cardiomyocyte cohesion via enhanced desmosomal assembly which is evidenced by enhanced DP/DSG2 localization at cell borders. It is the first step towards a novel therapeutic approach for arrhythmogenic cardiomyopathy.

## Introduction

The epidermal growth factor receptor (EGFR) is a ubiquitously expressed transmembrane receptor tyrosine kinase. Dysregulation of EGFR signaling often results in pathologies, such as different kinds of cancer, but also inflammatory skin and bowel disease or other skin, inflammatory or renal disorders (Bhargava et al., 2005, Kleinschmidt-DeMasters et al., 2005, Shia et al., 2005, da Cunha Santos et al., 2011, Campbell et al., 2014, Harskamp et al., 2016, Iwamoto et al., 2019). Today, EGFR inhibitors, either small inhibitory molecules (e.g., erlotinib, gefitinib or lapatinib) or inhibitory antibodies (e.g., cetuximab or panitumumab) are approved for the treatment of non-small cell lung cancer, pancreatic cancer, breast cancer, colorectal carcinoma and squamous cell carcinoma of head and neck (Westover et al., 2018, Sabbah et al., 2020). EGFR can be transactivated through G protein-coupled receptors via second messengers, several protein kinases or matrix metalloproteases (Noma et al., 2007, Makki et al., 2013). In addition, EGFR signaling can be activated, enhanced and mediated by the proto-oncogene tyrosine-protein kinase Src (SRC) (Chen et al., 2018). Vice versa, SRC can be activated by EGFR (Wilde et al., 1999). In the heart, EGFR was found to play a crucial role in cell metabolism, proliferation, cell survival, as well as hypertrophy and remodeling (Stroedecke et al., 2021). EGFR inhibition can lead to heart failure in the healthy heart, and a total knockout of the receptor can lead to severe hypertrophy and hypotension (Schreier et al., 2013, Kenigsberg et al., 2017, Guo et al., 2021). However, under pathological conditions, EGFR inhibition can be cardioprotective, as EGFR has been shown to be upregulated in a diabetes (Matrougui, 2010). Interestingly, EGFR inhibition protected vasculature and heart from diabetes-induced damages (Belmadani et al., 2008, Benter et al., 2009, Akhtar et al., 2012, Choi et al., 2012, El-Hashim et al., 2017). Furthermore, recent studies indicate that EGFR inhibition has a protective effect on cardiomyocytes under hypoxic conditions, i.e., after myocardial infarctions (Heliste et al., 2021). It seems that like in other tissues, EGFR signaling is tightly controlled in the heart, and due to its double role and the multitude of its downstream effectors, any alterations in EGFR signaling can have severe effects on heart function (Shraim et al., 2021, Forrester et al., 2016).

Arrhythmogenic cardiomyopathy (AC) is a disease most commonly caused by mutations in genes coding for desmosomal proteins, such as desmoglein 2 (DSG2), plakoglobin (PG), desmoplakin (DP), desmocollin 2 (DSC2), N-Cadherin (N-CAD) and plakophilin 2 (PKP2) (Gerull and Brodehl, 2020, Corrado et al., 2017). Hence, AC is considered as a hereditary disease of the desmosome. AC-causing mutations often lead to alterations in desmosomal contacts between cells, thereby weakening cellular cohesion. Several pathways regulating cardiomyocyte cohesion have been identified so far (Genet et al., 2012, Schlipp et al., 2014, Schinner et al., 2017, Zhao et al., 2019, Shoykhet et al., 2020). Furthermore, there is increasing evidence that desmosomes not only provide cellular cohesion, but also act as signaling hubs (Waschke, 2019, Muller et al., 2021). Thus, cardiac desmosomes are part of a complex signaling network, can influence, and are influenced by cellular signaling processes. Recent advancement in research suggests that similar molecular mechanisms are present in AC and pemphigus, an autoimmune skin disease, as well as inflammatory bowel diseases (Waschke, 2019, Bieber et al., 2020, Shoykhet et al., 2020, Schmitt and Waschke, 2021, Yeruva and Waschke, 2022).

In the context of pemphigus, EGFR inhibition led to increased cellular cohesion in keratinocytes (Walter et al., 2019), whereas in squamous carcinoma cell lines, EGFR inhibition increased protein levels of DSG2 and DSC2 (Lorch et al., 2004). However, EGFR inhibition by erlotinib in intestinal epithelial cells caused a decrease in cellular cohesion (Ungewiss et al., 2018), revealing that the role of EGFR can vary between different cells or tissues.

EGFR seems to play a major role in the heart function. Furthermore, links between EGFR and DSG2 and other desmosomal proteins have been established. Thus, in this study, we investigated whether inhibition of the EGFR signaling pathway can be used as a therapeutic option for AC, via regulating cardiomyocyte cohesion, using the *Jup*^-/-^ mouse model, which we characterized previously (Schinner et al., 2017, Schinner et al., 2020, Shoykhet et al., 2020). To this end, we used the well-known and clinically approved first-generation EGFR inhibitor erlotinib and an EGFR signaling molecule SRC inhibitor, PP2. Erlotinib was used to directly inhibit EGFR and PP2 to indirectly inhibit EGFR, by inhibiting the EGFR signaling pathway, since SRC acts both up- and downstream of EGFR. This approach was used as a proof of concept that EGFR inhibition, either using drug or a signaling inhibitor, rescues impaired cardiomyocyte cohesion.

## Results

### Inhibition of EGFR or SRC lead to positive adhesiotropy in HL-1 cardiomyocytes and murine cardiac slice cultures

In keratinocytes and intestinal epithelial cells, as well as in cancer cells, it is known that EGFR can modulate the desmosomal structure, assembly and signaling and vice versa (Lorch et al., 2004, Klessner et al., 2009, Bektas et al., 2013, Kamekura et al., 2014, Ungewiss et al., 2018). We used the cardiomyocyte-specific plakoglobin knockout (*Jup*^-/-^) mouse, representing a pathogenic AC model, which shows fibrotic replacement of myocardial tissue as well as arrhythmias (Schinner et al., 2020, Schinner et al., 2017, Shoykhet et al., 2020). In this model, we found increased EGFR protein expression in the *Jup*^-/-^ mice compared to wild type (*Jup*^+/+^) mice (Figure 1A and Supplementary Figure 1A). We next investigated the role of EGFR in cardiomyocyte cohesion. To this end, we inhibited EGFR directly using erlotinib or inhibited SRC using PP2, thereby inhibiting the EGFR pathway, and performed dispase-based dissociation assays with HL-1 cardiomyocytes and murine cardiac slice cultures. Therefore, treated HL-1 cell monolayers or murine cardiac slices were subjected to stress in order to challenge cell-cell contacts, upon which fragments of the monolayer (for HL-1 cells) or single cardiomyocytes (for murine cardiac slices) resulted.

**Figure 1.**
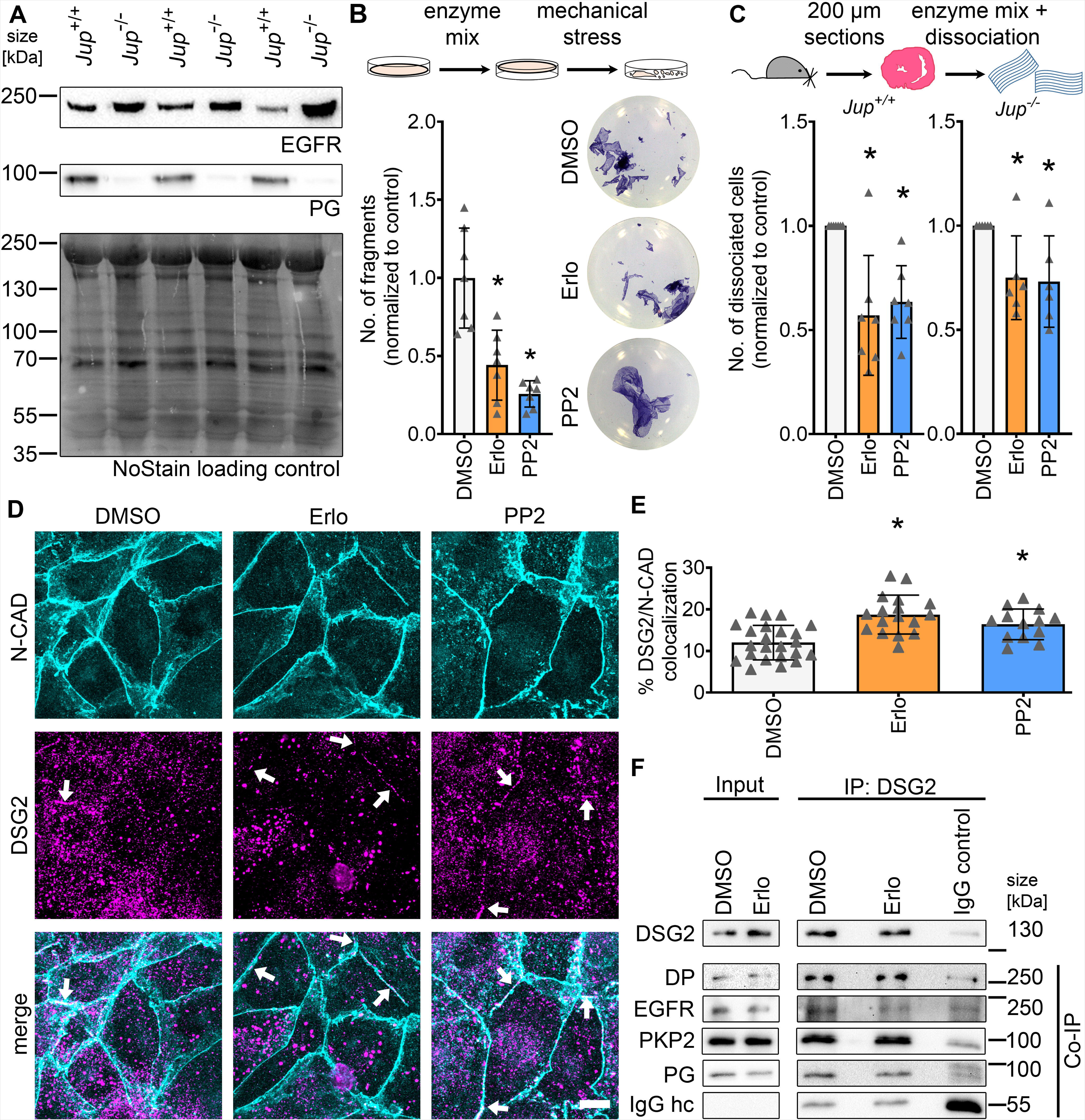
EGFR or SRC inhibition induces positive adhesiotropy in cardiomyocytes. **A**: Representative Western blot showing protein expression of EGFR and PG in *Jup*^+/+^ and *Jup*^-/-^ mice. **B**: Dispase-based dissociation assay in HL-1 cardiomyocytes upon inhibition of EGFR or SRC by erlotinib or PP2, respectively, with representative pictures of the wells, * p ≤ 0.05 (p=0.0003 for erlotinib and p<0.0001 for PP2), 1-way ANOVA with Holm-Sidak correction, N=7 biological replicates. **C**: Dispase-based dissociation assay in murine cardiac slice cultures obtained from *Jup*^+/+^ and *Jup*^-/-^ mice upon inhibition of EGFR or SRC, * p ≤ 0.05 (*Jup*^+/+^ mice: p=0.0012 for DMSO vs erlotinib and p=0.0024 for DMSO vs PP2. *Jup*^-/-^ mice: p=0.0322 for DMSO vs erlotinib and PP2), 1-way ANOVA with Holm-Sidak correction, N=7 for *Jup*^+/+^ mice and N=6 for *Jup*^-/-^ mice. **D**: Maximum projections of immunostainings in HL-1 cardiomyocytes for N-CAD and DSG2 showing an increase of DSG2 at the cell borders after erlotinib or PP2 treatments. Z-scans spanning the whole cell volume, z-steps = 0.25 µm. White arrows indicate areas of increased DSG2 localization at the cell membrane. Scale bar: 10 µm. **E:** Quantification of colocalization of N-CAD and DSG2 in HL-1 cardiomyocytes, * p ≤ 0.05 (p<0.0001 for DMSO vs erlotinib and p=0.0042 for DMSO vs PP2), 1-way ANOVA with Holm-Sidak correction. Each data point represents one individual cell border, N=5 biological replicates. **F:** Representative Western blots for immunoprecipitation of DSG2, co-immunoprecipitation of DP, EGFR, PKP2 and PG, IgG heavy chain (IgG hc) served as loading control for immunoprecipitated samples, N=3 biological replicates.

We show that both EGFR or SRC inhibition enhanced cardiomyocyte cohesion, which we term as positive adhesiotropy, in HL-1 cardiomyocytes and murine cardiac slice cultures (Figure 1B and 1C). Inhibition of EGFR or SRC was confirmed by assessing the phosphorylation levels of their downstream targets: ERK and EGFR (Supplementary Figure 1B-D). We also performed immunostaining of N-CAD and DSG2 in HL-1 cardiomyocytes and document an increase of DSG2 at the cell borders after EGFR or SRC inhibition, compared to controls (Figures 1D and 1E). Next, to assess whether EGFR is in a complex with desmosomal proteins, we immunoprecipitated DSG2 from HL-1 lysates and performed immunoblots for EGFR, PG, DP and PKP2. Indeed, EGFR was found in complex with DSG2 in addition to PG, DP and PKP2 (Figure 1F). Nevertheless, treatment with erlotinib did not alter the composition of this complex.

These experiments showed that inhibition of either EGFR or SRC leads to positive adhesiotropy in HL-1 cardiomyocytes and murine cardiac slice cultures, paralleled by enhanced DSG2 localization along cell junctions.

### Inhibition of EGFR increases the DSG2 binding event frequency at cell borders

Since we observed an increase in DSG2 localization at the membrane after EGFR or SRC inhibition, we assessed the role of DSG2 in strengthening cellular cohesion upon EGFR inhibition in cardiomyocytes. For this, atomic force microscopy (AFM) measurements with HL-1 cardiomyocytes were used. Here, a cantilever with a protein-coated tip approached the cells, where the protein at the tip could eventually interact with proteins at the cell surface. We measured the binding frequency of DSG2, DSC2, or N-CAD-coated tips, as well as the unbinding forces at cell borders and cell surfaces. We observed an increase in binding frequency at cell borders for DSG2-coated tips only (Figure 2A). In the case of N-CAD coated tips no change in binding frequency was observed (Figure 2B). However, in the case of DSC2-coated tips, reduced binding frequency at the cell surface, but not the cell border, was found (Figure 2C). On the other hand, no changes in unbinding forces for any of the proteins analyzed were observed (Figure 2D-F). These data indicate that enhanced binding frequency of DSG2 correlates with the strengthening of intercellular adhesion in HL-1 cardiomyocytes upon EGFR inhibition.

**Figure 2.**
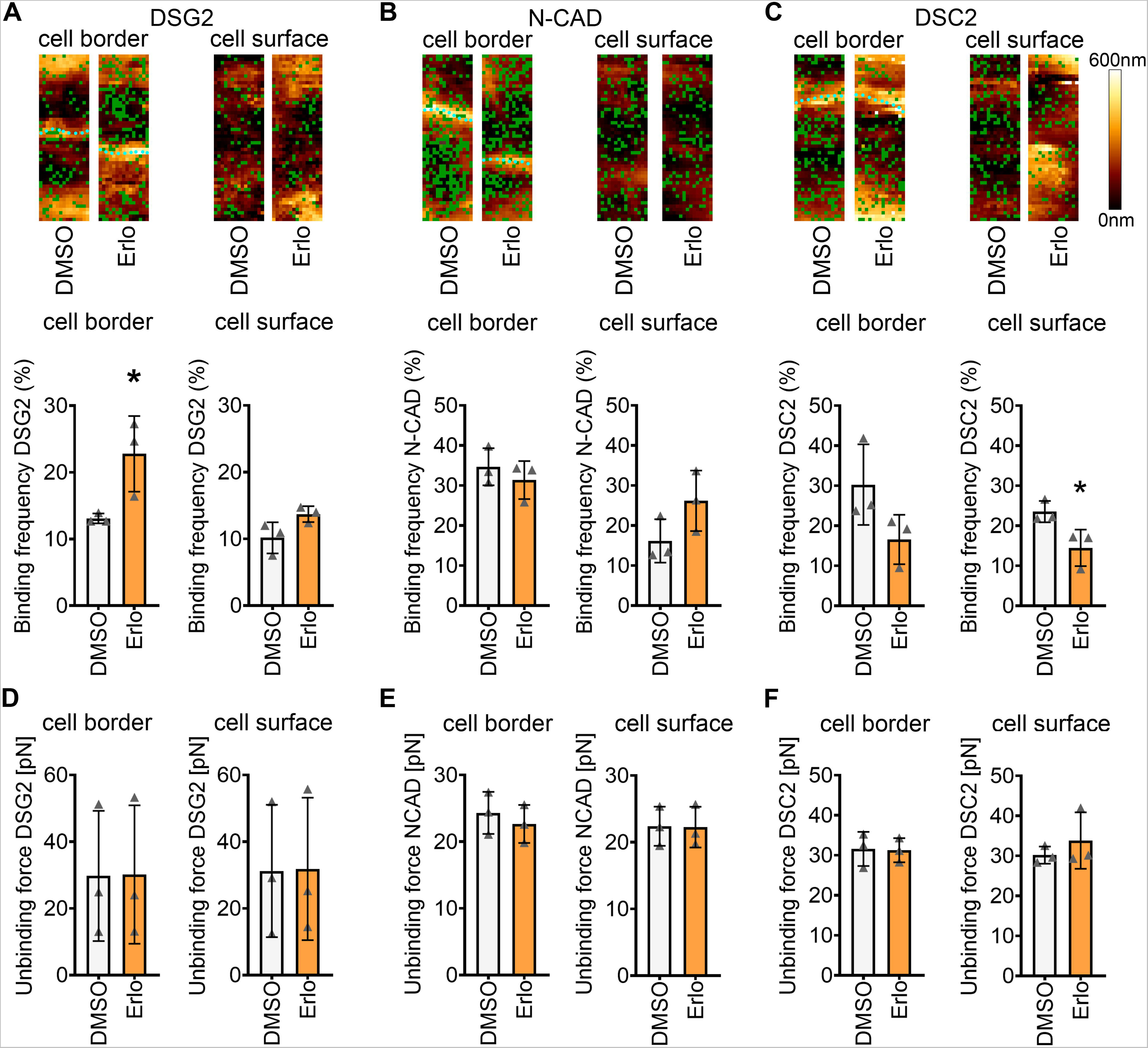
EGFR inhibition leads to increased DSG2 binding frequency at cell borders. **A-C**: Binding frequency and topography during atomic force microscopy (AFM) measurements on HL-1 cardiomyocytes before and after erlotinib treatment at or close to cell borders and cell surfaces with tips coated with **A**: DSG2, **B**: N-CAD, **C**: DSC2. Measurements were first performed in 90 minutes DMSO-treated samples, then medium was changed, erlotinib was added and experiment was performed after 90 minutes. Green dots in topography images indicate binding events; cyan dots indicate cell borders. Topography images are 1.5 µm x 5 µm. In the graphs, per data-point, 1500 curves were analyzed across two areas (1.5 µm x 5 µm). * p ≤ 0.05. Statistical significance was calculated between DMSO vs erlotinib. (DSG2: cell borders p=0.0426, cell surface: p=0.0785; N-CAD: cell borders p=0.433, cell surface: p=0.1357; DSC2: cell borders p=0.1151, cell surface: p=0.041), unpaired Student’s *t*-test, N=3 biological replicates. **D-F**: Unbinding forces measured during AFM measurements on HL-1 cardiomyocytes at cell borders and cell surfaces with tips coated with **D**: DSG2, **E**: N-CAD, **f**: DSC2. Per data-point, 1500 curves were analyzed across two areas (1.5 µm x 5 µm). * p ≤ 0.05. Statistical significance was calculated between DMSO vs erlotinib. (DSG2 cell borders 0.9808, cell surface p=0.9722; N-CAD: p=0.5380, cell surface p=0.958; DSC2: p=0.9258, cell surface p=0.4421), unpaired, Student’s *t*-test, N=3 biological replicates.

### Apart from DSG2, positive adhesiotropy induced by EGFR or SRC inhibition is also dependent on DP

To further understand the mechanism of erlotinib or PP2-induced positive adhesiotropy, we performed siRNA-mediated knockdown in HL-1 cardiomyocytes. Firstly, we knocked down *Egfr*, which did not alter cellular cohesion (Figure 3A and Supplementary Figure 2A). We then knocked down *Egfr* and treated the cells with erlotinib or PP2. Upon *Egfr* knockdown, erlotinib-induced positive adhesiotropy was abolished, confirming that the effect observed after erlotinib treatment is mediated through EGFR. Interestingly, SRC inhibition was still effective after *Egfr* knockdown, indicating that the PP2 effect is either independent of EGFR or that SRC is downstream of EGFR (Figure 3B and Supplementary Figure 2A). Next, we knocked down *Dsp* or *Dsg2* (Figure 3C). Knocking down *Dsp* or *Dsg2* did not alter cellular cohesion (Figure 3D and Supplementary Figure 2B and 2C), as observed before (Shoykhet et al., 2020, Schinner et al., 2020). Both, erlotinib or PP2 enhanced cellular cohesion after *Dsg2* knockdown. This suggests that the remaining DSG2 protein might be sufficient enough to mediate the positive adhesiotropic effect of EGFR or SRC inhibition. In previous reports inhibition of EGFR enhanced DP expression in breast cancer cells and led to enhanced DP at the cell border in oral squamous cell carcinoma cells (Lorch et al., 2004, Pang et al., 2004). Based on these observations, we knocked down *Dsp*. EGFR or SRC inhibition-mediated positive adhesiotropy was abrogated upon *Dsp* knockdown, indicating the crucial role of DP in erlotinib or PP2-mediated positive adhesiotropy (Figure 3E and Supplementary Figure 2C). These data highlight the role of DP, along with DSG2, in positive adhesiotropy induced by EGFR or SRC inhibition.

**Figure 3.**
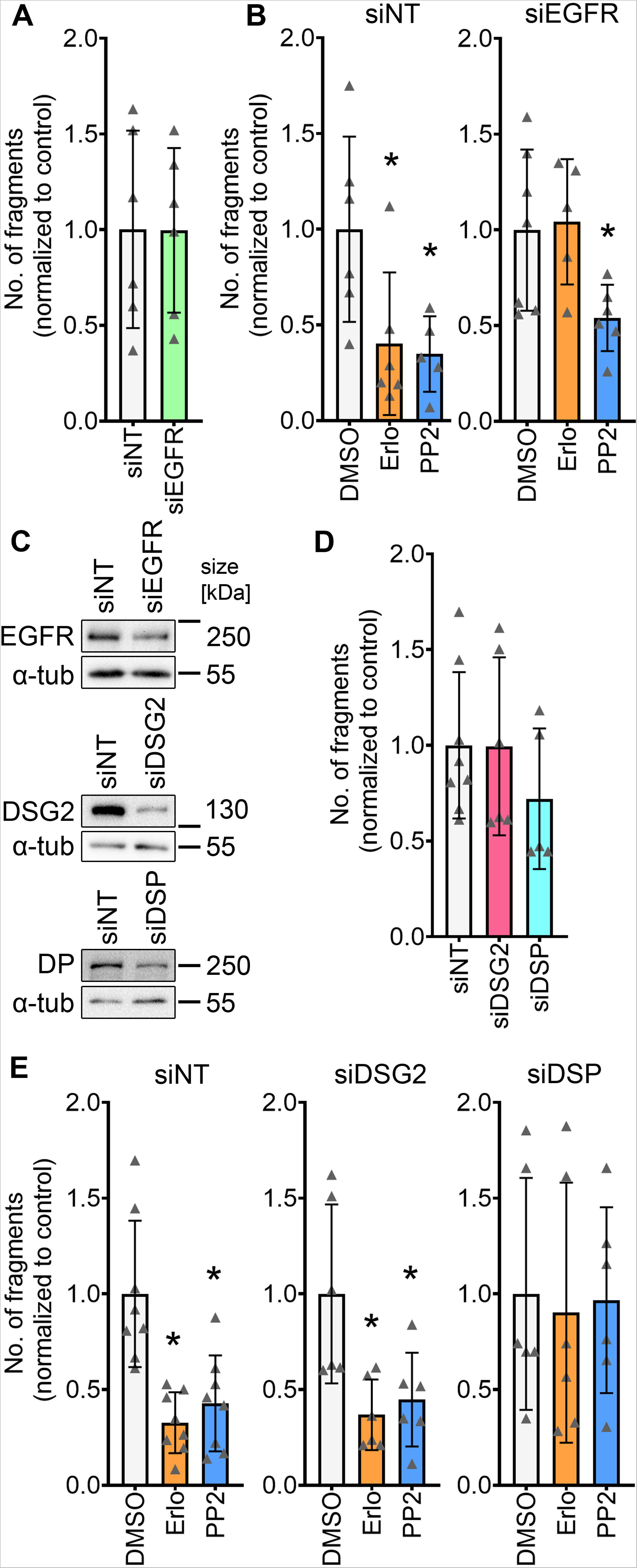
Positive adhesiotropy induced by EGFR or SRC inhibition is dependent on DP. **A**: Dispase-based dissociation assay in HL-1 cardiomyocytes after siRNA-mediated knockdown of EGFR and non-target (NT) control knockdown. Unpaired Student’s *t*-test (p=0.9858), N=6 biological replicates. **B**: Dispase-based dissociation assay in HL-1 cardiomyocytes after siRNA-mediated knockdown of EGFR and treatments with erlotinib or PP2. *p ≤ 0.05 (siNT: p=0.0264 for DMSO vs erlotinib and PP2; siEGFR: p=0.8257 for DMSO vs erlotinib and p=0.0494 for DMSO vs PP2), 1-way ANOVA with Holm-Sidak correction, N=6. **C**: Representative Western blot confirmation of siRNA-mediated knockdown efficiency for EGFR, DP and DSG2, α-tubulin served as a loading control, N=6 biological replicates. **D**: Dispase-based dissociation assay in HL-1 cardiomyocytes after siRNA-mediated knockdown of DSG2 or DP *p ≤ 0.05 (p=0.982 for siNT vs siDSG2 and p=0.2204 for siNT vs siDSP), unpaired Student’s *t*-test. **E**: Dispase-based dissociation assay in HL-1 cardiomyocytes after siRNA-mediated knockdown of DSG2 or DP and treatments with erlotinib or PP2. *p ≤ 0.05 (siNT: p=0.0002 for DMSO vs erlotinib and p=0.001 for DMSO vs PP2; siDSG2: p=0.0081 for DMSO vs erlotinib and p=0.0097 for DMSO vs PP2; siDSP: p=0.9512 for DMSO vs erlotinib and PP2), 1-way ANOVA with Holm-Sidak correction, N=6-8 biological replicates.

### EGFR or SRC inhibition leads to increased area composita length in HL-1 cardiomyocytes

Since both, DSG2 and DP seem to be involved in the positive adhesiotropy induced by EGFR or SRC inhibition, we next performed immunostaining of these proteins together with Phalloidin to stain the cortical actin cytoskeleton. Confocal microscopy images revealed an increase in the localization of DP and DSG2 at cell borders after inhibition of EGFR or SRC (Figure 4A-C). At the same time DSG2 and DP staining looked enlarged in erlotinib and PP2-treated cells compared to vehicle controls, which indicated changes at the area composita, that were shown to be present in intercellular junctions of HL-1 cells (Schlipp et al., 2014). Therefore, we performed stimulated emission depletion (STED) imaging for DSG2 and DP, which showed that area composita number remained unchanged, whereas the area composita length increased after EGFR or SRC inhibition (Figure 4D-F). Apart from that, we observed an increase in desmin (DES) insertion into areae compositae upon treatment with erlotinib or PP2 (Supplementary Figure 3).

**Figure 4.**
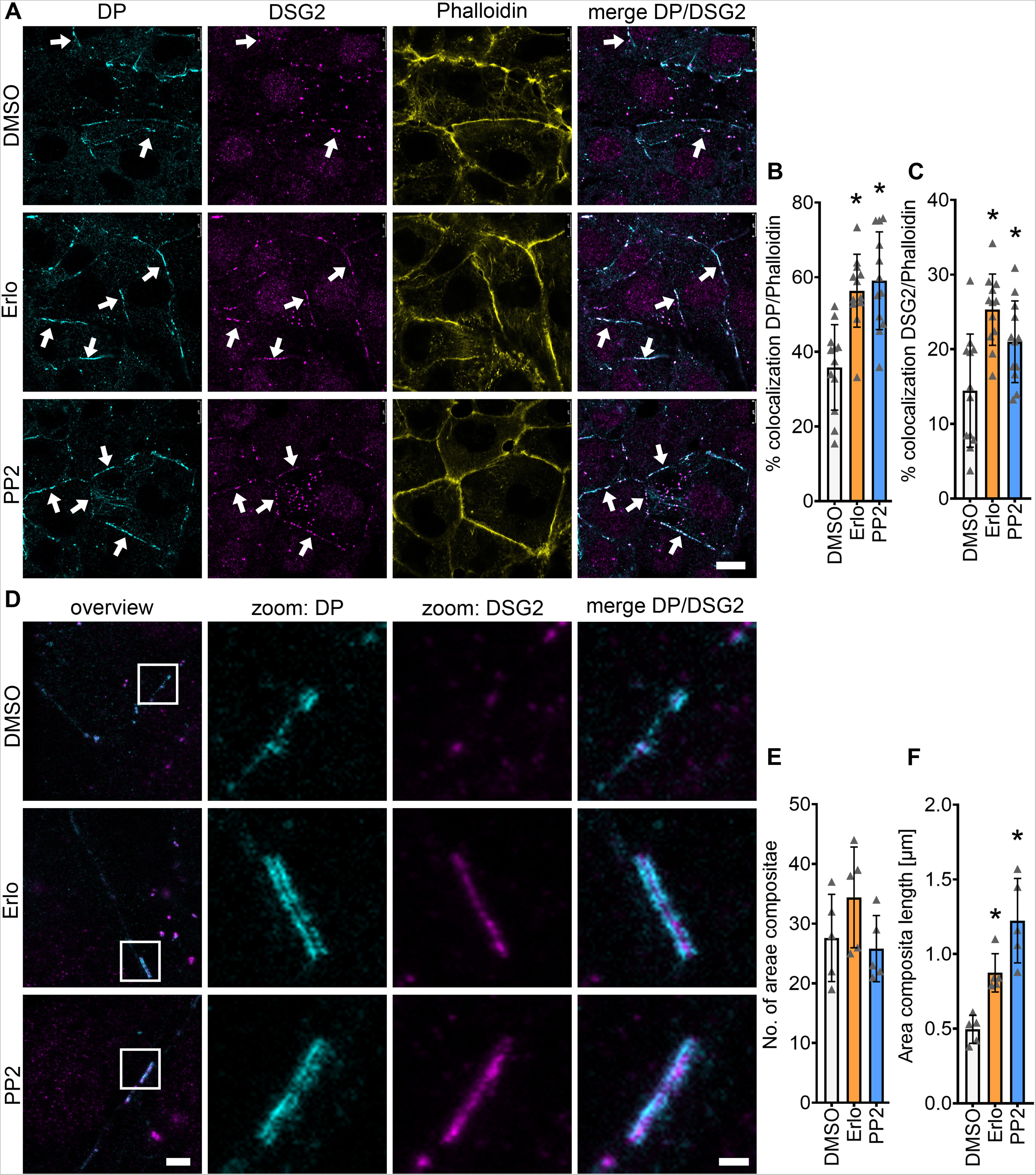
Increased DP and DSG2 recruitment to the cell membranes leads to longer areae compositae upon EGFR or SRC inhibition. **A**: Immunostaining of DP and DSG2 in HL-1 cardiomyocytes with Phalloidin as membrane marker. Scale bar: 10 µm. White arrows represent an increase in DSG2 and DP localization to the membrane compared to DMSO. **B**: Quantification of colocalization of DP and Phalloidin. *p ≤ 0.05 (p=0.0001 for DMSO vs erlotinib and p<0.0001 for DMSO vs PP2), 1-way ANOVA with Holm-Sidak correction. **C**: Quantification of colocalization of DSG2 and Phalloidin. *p ≤ 0.05 (p=0.0002 for DMSO vs erlotinib and p=0.0122 for DMSO vs PP2), 1-way ANOVA with Holm-Sidak correction. Each data point represents one individual cell border, N=6 biological replicates. **D**: STED images of HL-1 cardiomyocytes stained for DP and DSG2. Scale bar in overviews: 2 µm, scale bar in zooms: 500 nm. **E**: Number of areae compositae over six areas (15 µm x 15 µm) from N=5 biological replicates, *p ≤ 0.05 (p=0.2959 for DMSO vs erlotinib and p=0.6994 for DMSO vs PP2), 1-way ANOVA with Holm-Sidak correction. **F**: Quantification of area composita length. *p ≤ 0.05 (p=0.0078 for DMSO vs erlotinib and p=0.0001 for DMSO vs PP2), 1-way ANOVA with Holm-Sidak correction, N=4 biological replicates.

### Enhanced DP and DSG2 staining in murine cardiac slices after EGFR or SRC inhibition

To confirm the findings from HL-1 cardiomyocytes in a mouse model, we performed immunostaining of murine cardiac slice cultures, both in *Jup^+/+^* and in *Jup*^-/-^ mice. As observed before (Yeruva et al., 2020), DSG2 was less expressed in the ICDs of *Jup*^-*/-*^ mice compared to *Jup*^+/+^ mice (Figure 5A and D). No changes in staining intensity or ICD lengths between mediator-treated and untreated slice cultures were found in murine cardiac slice cultures from *Jup*^+/+^ or *Jup*^-/-^ mice. Nonetheless, both DP and DSG2 staining were broader after EGFR or SRC inhibition (Figure 5A-C). Interestingly, in cardiac slices obtained from *Jup*^-/-^ mice, only DSG2 staining width was increased after EGFR or SRC inhibition, whereas changes in DP staining width were not observed (Figure 5D-F). Taken together, immunostaining showed increased DSG2 and DP localization at cellular junctions after EGFR or SRC inhibition.

**Figure 5.**
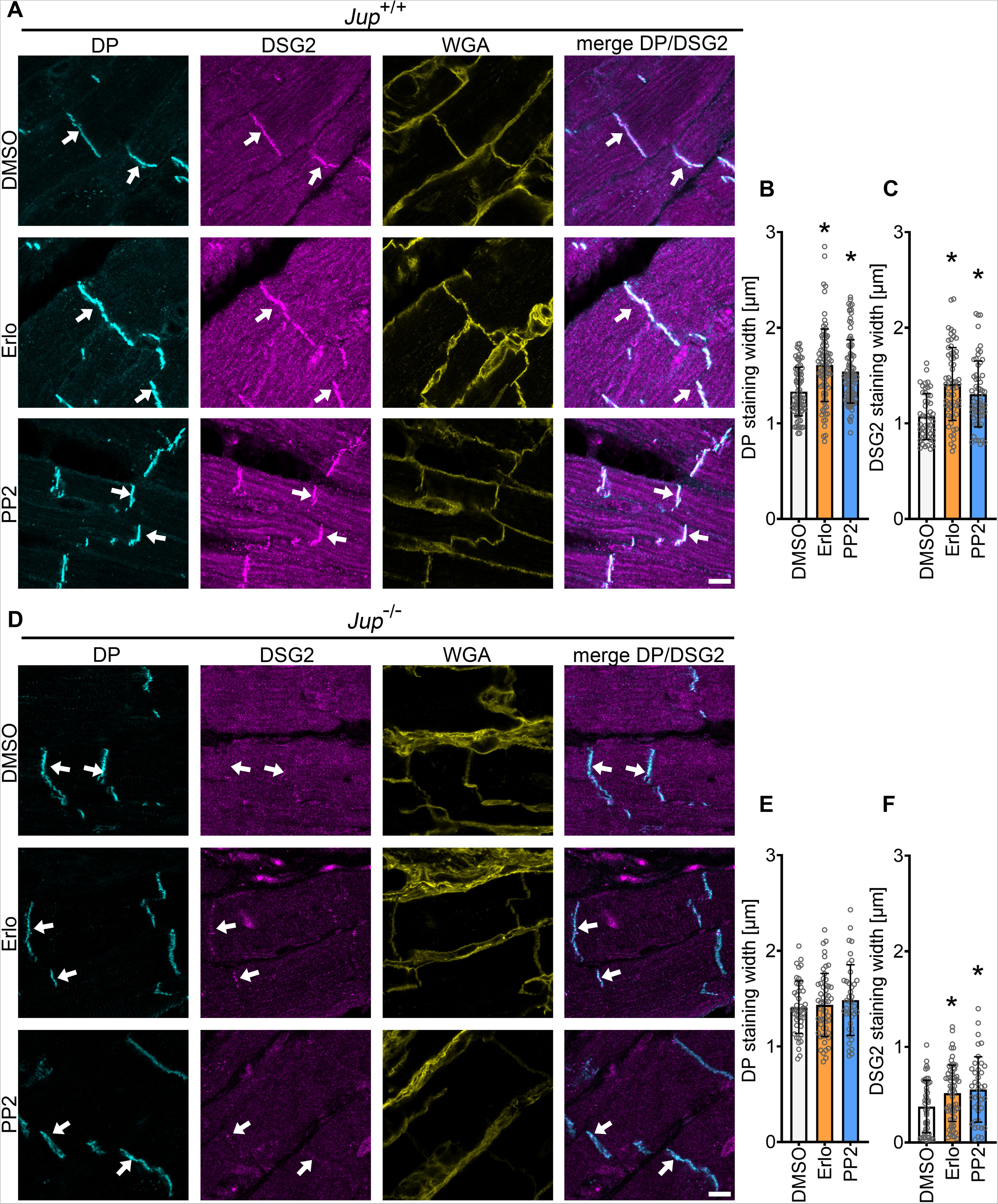
EGFR or SRC inhibition leads to increased recruitment of DP and DSG2 into the ICDs in *Jup*^+/+^ and *Jup*^-/-^ mice. **A**: Immunostaining of DP and DSG2 in *Jup*^+/+^ murine cardiac slices with WGA as membrane marker. Scale bar: 8 µm. **B**: DP staining width. **C**: DSG2 staining width. * p ≤ 0.05, (DP staining width: p<0.0001 for DMSO vs erlotinib and PP2; DSG2 staining width: p<0.0001 for DMSO vs erlotinib and p=0.0004 for DMSO vs PP2), 1-way ANOVA with Holm-Sidak correction. Each data point represents one ICD, N=4 mice. **D**: Immunostainings of *Jup*^-/-^ murine cardiac slices for DP and DSG2 with WGA as membrane marker. Scale bar: 8 µm. **E**: DP staining width. **F**: DSG2 staining width. * p ≤ 0.05 (DP staining width: p=0.7121 for DMSO vs erlotinib and p=0.5042 for DMSO vs PP2; DSG2 staining width: p=0.0189 for DMSO vs erlotinib and p=0.0147 for DMSO vs PP2), 1-way ANOVA with Holm-Sidak correction. Each data point represents one ICD, N=4 mice. White arrows represent an increase staining width in DSG2 or DP staining compared to DMSO.

### EGFR or SRC inhibition promotes desmosomal assembly in HL-1 cardiomyocytes

Since the localization of DSG2 at cell borders was increased after EGFR or SRC inhibition, we investigated whether EGFR inhibition alters DSG2 mobility. To address this, we transfected HL-1 cardiomyocytes with a DSG2-GFP construct and performed fluorescence recovery after photobleach (FRAP) experiments. We neither found changes in the halftime of recovery (τ), nor in the immobile fraction after EGFR inhibition (Figures 6A and 6B). Furthermore, we did not observe significant protein localization changes between cytosolic and membrane-bound fractions in Triton-X-100 assays (Supplementary Figure 4A and 4B). Therefore, we next studied desmosomal assembly and performed a Ca^2+^-switch assay by depleting cells of Ca^2+^ for 90 minutes, using the Ca^2+^-chelator EGTA to disrupt desmosomes. We performed dispase-based dissociation assays after changing the medium to Ca^2+^- containing medium together with erlotinib or PP2. EGFR or SRC inhibition led to increased cellular cohesion after a Ca^2+^-switch, indicating increased desmosomal assembly (Figure 6C and Supplementary Figure 4C). Indeed, immunostaining for DP and DSG2 revealed that both EGFR and SRC inhibition led to enhanced recruitment of DP and DSG2 to the cell borders after the Ca^2+^-switch in HL-1 cardiomyocytes (Figure 6D-F). Immunoprecipitation of DP after EGFR and SRC inhibition revealed more association of PG and DP in HL-1 cardiomyocytes, while PKP2 and DSG2 association with DP was only increased upon SRC inhibition (Figure 6G). These data confirmed that EGFR and SRC inhibition led to enhanced desmosomal assembly.

**Figure 6.**
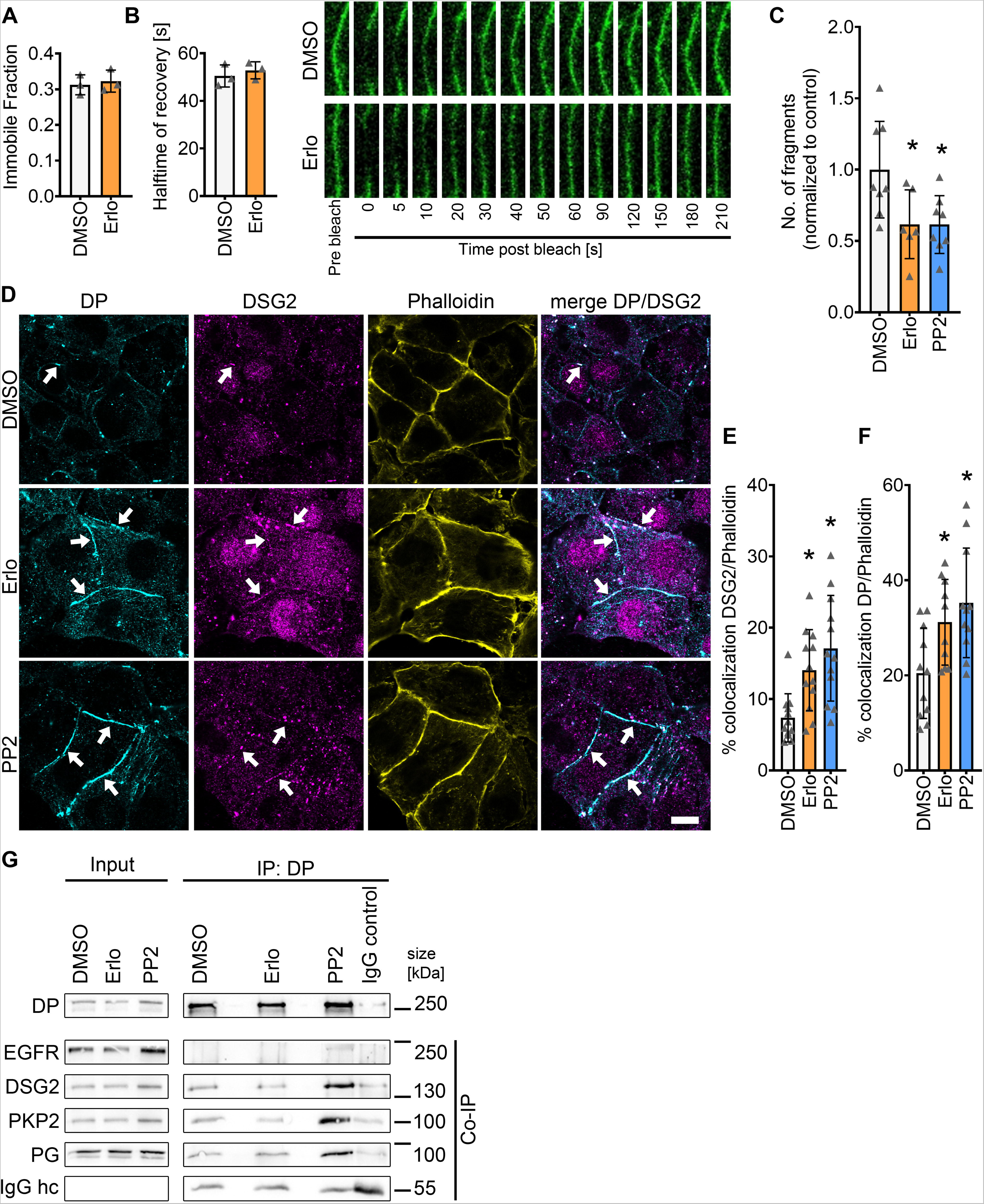
Increased recruitment of DP and DSG2 towards the cell membrane is achieved by enhanced desmosome assembly. **A-B**: Fluorescence recovery after photobleach (FRAP) measurements in HL-1 cardiomyocytes transfected with DSG2-GFP and treated with DMSO or erlotinib. Measurements were performed 60-120 minutes post treatment **A**: Immobile fraction **B**: Halftime of recovery (τ). * p ≤ 0.05 (immobile fraction p=0.6933; τ p=0.5252 for DMSO vs erlotinib), unpaired Student’s *t*-test, N=4 biological replicates. **C:** Dispase-based dissociation assay in HL-1 cardiomyocytes after 90 minutes of Ca^2+^-depletion and subsequent Ca^2+^- repletion together with erlotinib or PP2. *p ≤ 0.05 (p=0.0201 for DMSO vs erlotinib and PP2), 1-way ANOVA with Holm-Sidak correction, N=6 biological replicates. **D:** Immunostaining of DP and DSG2 in HL-1 cardiomyocytes with Phalloidin as membrane marker after 90 minutes of Ca^2+^-depletion and subsequent Ca^2+^-repletion together with erlotinib or PP2. White arrows indicate areas of increased DP or DSG2 recruitment to the cell membrane. Scale bar: 10 µm. **E-F:** Quantification of colocalization of **E**: DP or **F**: DSG2 and Phalloidin, * p ≤ 0.05 (DP: p=0.0216 for DMSO vs erlotinib and p=0.0036 for DMSO vs PP2; DSG2: p=0.0073 for DMSO vs erlotinib and p=0.0004 for DMSO vs PP2), 1-way ANOVA with Holm-Sidak correction, N=4 biological replicates. Each data point represents one individual cell border. **G:** Immunoprecipitation of DP from HL-1 lysates, co-immunoprecipitation of EGFR, DSG2, PKP2 and PG, N=3 biological replicates.

### Positive adhesiotropy induced by EGFR inhibition is mediated by ROCK

We next aimed to unravel potential mechanisms of EGFR-inhibition mediated positive adhesiotropy. Since the EGFR signaling cascade involves several different kinases, we performed a multiplex kinase activity array (PamGene Kinase) utilizing HL-1 cardiomyocytes. Here we compared kinase activity in erlotinib-treated samples to control samples after 15, 30 and 60 minutes of incubation (Table S1). After 30 minutes of EGFR inhibition, we observed an increase in PCTAIRE2, also known as cyclin dependent kinase 17 (CDK17), Rho associated kinase 1 (ROCK1) and protein kinase N1 (PKN1) activity. Since Rho family members are known to be involved in cellular cohesion (Kaibuchi et al., 1999, Fukata et al., 1999, Evers et al., 2000), and PKN1 is an effector kinase of RhoA (Amano et al., 1996, Flynn et al., 1998), which is upstream of ROCK, we focused on ROCK. Indeed, inhibition of ROCK by Y27632 decreased cellular cohesion in HL-1 cardiomyocytes and prevented erlotinib-induced positive adhesiotropy in HL-1 cardiomyocytes (Figure 7A), indicating that ROCK is involved in cardiomyocyte cohesion. We then performed immunostainings and found that upon ROCK inhibition, erlotinib did not enhance DP and DSG2 staining at the cell borders (Figures 7B-D). To further characterize whether ROCK is involved in desmosome assembly, we performed dispase-based dissociation assays after a Ca^2+^-switch in HL-1 cardiomyocytes. ROCK inhibition alone did not decrease cellular cohesion after a Ca^2+^-switch, whereas EGFR inhibition-mediated positive adhesiotropy was still prevented (Figure 7E).

**Figure 7.**
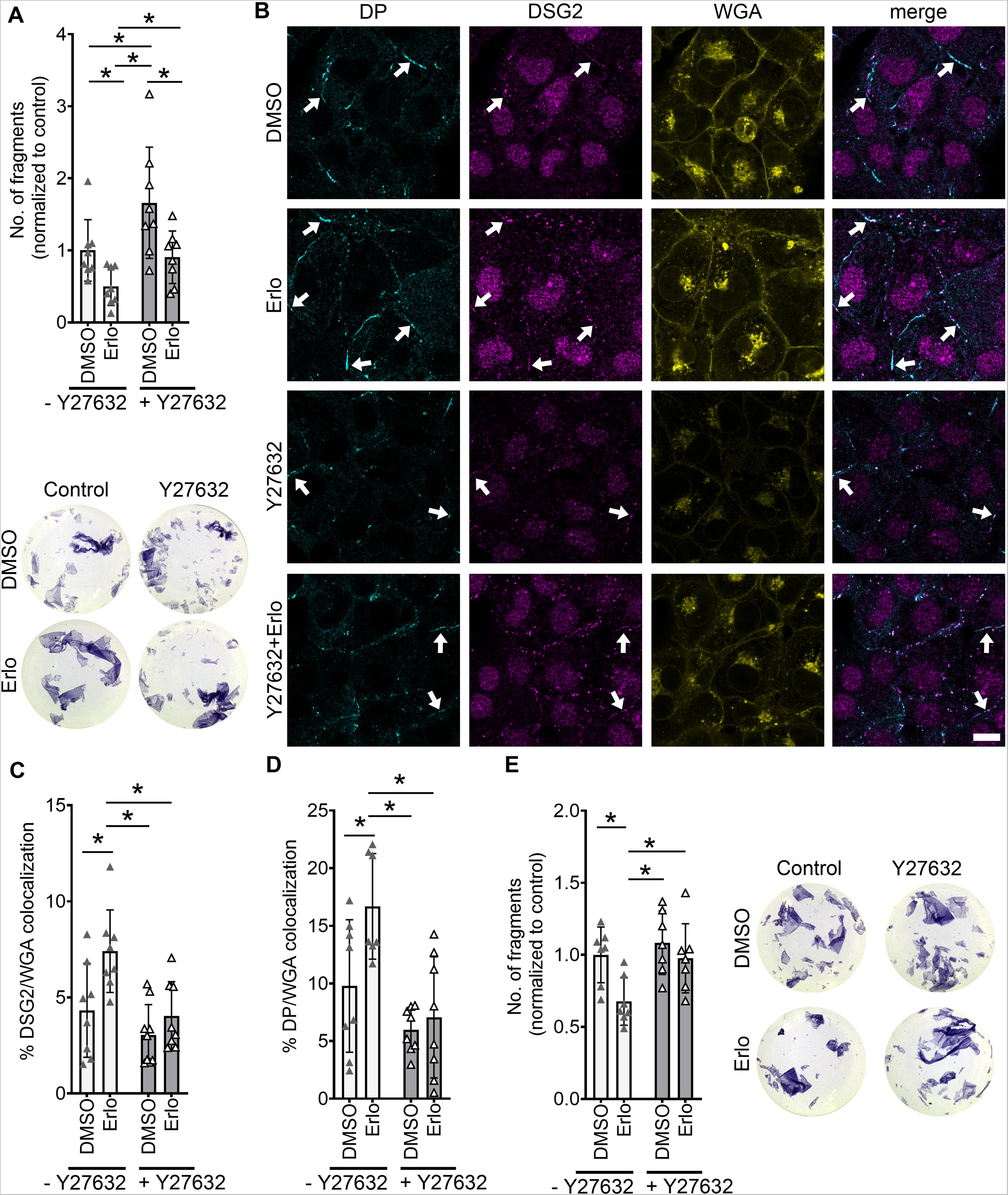
Erlotinib enhanced cardiomyocyte cohesion and desmosomal assembly are mediated by ROCK in HL-1 cardiomyocytes. **A:** Dispase-based dissociation assay in HL-1 cardiomyocytes, after treatment with erlotinib with and without Y27632. Y27632 was added 30 minutes prior to erlotinib incubation for 60 minutes, with representative pictures of the wells, (p=0.0095 for control vs Y27632; p=0.0287 for control vs erlotinib; p=0.5265 for control vs Y27632+erlotinib; p=0.0005 for Y27632 vs erlotinib; p=0.0056 for Y27632 vs Y27632+erlotinib; p=0.0482 for erlotinib vs Y27632+erlotinib), N=8 biological replicates, 2-Way ANOVA with Holm Sidak’s multiple comparison test. **B**: Immunostaining of DP and DSG2 in HL-1 cardiomyocytes with WGA as membrane marker after erlotinib treatment with and without Y26732, as in A. White arrows indicate areas of increased DP or DSG2 recruitment to the cell membrane. Scale bar: 10 µm. **C-D:** Quantification of colocalization of **C**: DP or **D**: DSG2 and WGA, 2-Way ANOVA with Holm-Sidak’s multiple comparison test * p ≤ 0.05 (DP: p=0.300 for Control vs Y27632, p= 0.0299 for Control vs erlotinib, p=0.4374 for Control vs Y27632+erlotinib, p=0.0008 for Y27632 vs erlotinib, p=0.6444 for Y27632 vs Y23632+erlotinib, p=0.0021 for erlotinib vs Y27632+erlotinib; DSG2: p=0.4857 for Control vs Y27632, p=0.0181 for Control vs erlotinib, p=0.7723 for Control vs Y27632+erlotinib, p=0.0006 for Y27632 vs erlotinib, p=0.5359 for Y27632 vs Y27632+erlotinib, p=0.0107 for erlotinib vs Y27632+erlotinib), N=3 biological replicates. Each data point represents one individual cell border. **E:** Dispase-based dissociation assay in HL-1 cardiomyocytes after 90 minutes of Ca^2+^-depletion and treatment with erlotinib with and without Y26732 with representative pictures of the wells. 2-Way ANOVA with Holm-Sidak’s multiple comparison test * p ≤ 0.05 (p=0.7069 for Control vs Y27632, p=0.0368 for Control vs erlotinib, p=0.8279 for Control vs Y27632+erlotinib, p=0.0070 for Y27632 vs erlotinib, p=0.7069 for Y27632 vs Y23632+erlotinib, p=0.0489 for erlotinib vs Y27632+erlotinib), N=7 biological replicates.

Taken together, these data suggest that the EGFR inhibition-induced positive adhesiotropy and enhanced desmosomal assembly is mediated by ROCK.

## Discussion

AC is a disease with complex pathogenesis, which at least in part is caused by disturbed desmosome turnover (Liang et al., 2021). The current therapeutic options for AC patients include changes in lifestyle, treatment with antiarrhythmic drugs, catheter ablation, implantable cardiac defibrillator, and end-stage heart transplantation (Corrado et al., 2017). However, in patients carrying mutations in genes coding for proteins of the desmosomal complex, lifestyle changes, such as restraining from physical endurance activities and β-blocker therapy are most commonly used (Corrado et al., 2017). In our previous work, we discovered adrenergic signaling via PKA-mediated PG-phosphorylation at serine 665 as a driving force for DSG2-mediated enhanced cardiomyocyte cohesion, which we termed positive adhesiotropy (Schlipp et al., 2014, Schinner et al., 2017, Shoykhet et al., 2020). This indicated that cardiomyocyte adhesion at ICDs is precisely regulated and can be enhanced via drugs. Because EGFR inhibitors are used in clinics for the treatment of several cancers (Westover et al., 2018, Sabbah et al., 2020), in this work, we characterized the role of EGFR on cardiomyocyte cohesion and the efficacy of erlotinib to modulate adhesion. Indeed, we show that inhibition of EGFR and its effector molecule, SRC, in HL-1 cardiomyocytes and murine cardiac slice cultures from *Jup*^+/+^ or *Jup*^-/-^ mice enhanced cardiomyocyte cohesion, paralleled by increased desmosome assembly into the area composita (Figure 8).

**Figure 8.**
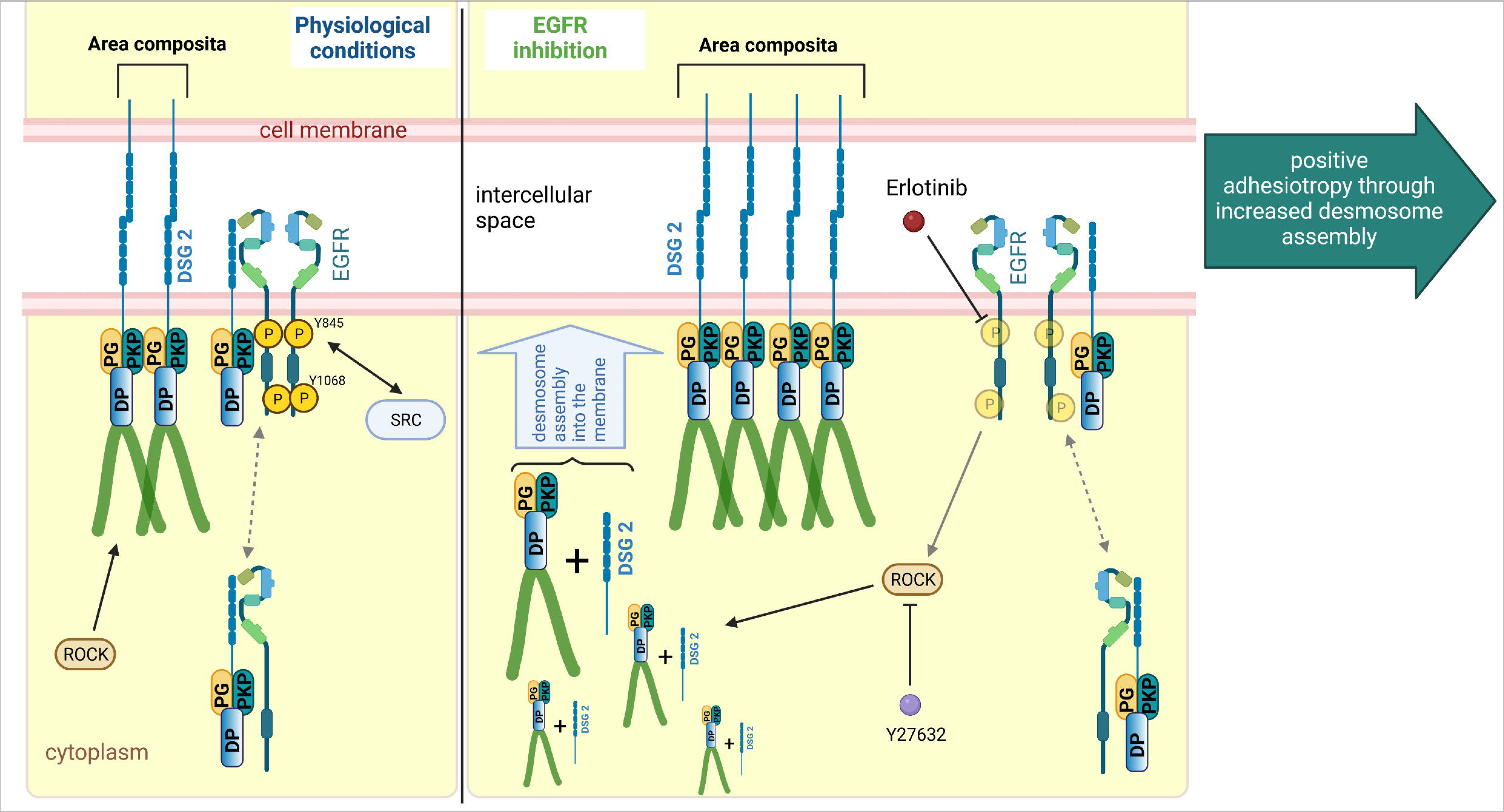
Schematic overview of EGFR inhibition-mediated positive adhesiotropy. EGFR exists in complex with DSG2, along with DP, PG and PKP2, either at the cell borders or in the cytoplasm. Under physiological conditions ROCK is necessary for basal cardiomyocyte cohesion. EGFR inhibition by erlotinib further elevates ROCK that leads to positive adhesiotropy via enhanced desmosome assembly and area composita length. Inhibition of ROCK completely abrogates erlotinib-mediated positive adhesiotropy.

As a first step, we analyzed the expression of EGFR in *Jup*^+/+^ or *Jup*^-/-^ mice hearts and found an increase in EGFR expression in the hearts of *Jup*^-/-^ mice. To our knowledge, this is the first instant of finding EGFR overexpression in murine models of AC. In fact, EGFR activation and signaling was shown to promote TGF-β dependent renal fibrosis (Chen et al., 2012, Samarakoon et al., 2013). Since in AC, replacement of myocardium with fibrotic tissue is a common phenomenon (Corrado et al., 2017), we believe that the observed EGFR overexpression might be related to fibrosis observed in *Jup*^-/-^ mice (Shoykhet et al., 2020, Yeruva et al., 2020, Schinner et al., 2017). It has been shown that upon treatment with the EGFR inhibitor gefitinib, inflammation and hypertrophy caused by chronic treatment with the adrenergic mediator isoprenaline, was ameliorated in mice (Grisanti et al., 2015, Talarico et al., 2014). Moreover, increased EGFR expression was positively correlated with the size of atherosclerotic plaques (Lamb et al., 2004). However, mice with a strong downregulation of EGFR in the heart, have severe hypertrophy and hypotension (Schreier et al., 2013).

We used erlotinib or PP2 to inhibit EGFR or SRC kinases, respectively, and focused more on erlotinib, since it is a clinically approved drug. Dispase-based dissociation assays using erlotinib or PP2 in HL-1 cardiomyocytes and murine cardiac slice cultures from *Jup*^+/+^ or *Jup*^-/-^ mice revealed that both inhibitors enhance cardiomyocyte cohesion. Similar to our findings in this study, EGFR inhibition has been shown to increase cellular cohesion in squamous cell carcinomas (Lorch et al., 2004), whereas in keratinocytes EGFR inhibition protected against DSG3 autoantibody-induced loss of keratinocyte adhesion (Walter et al., 2019). Immunoprecipitation revealed EGFR along with PKP2, DP and PG being in a complex with DSG2. In intestinal epithelial cells, as well as in a cell-free AFM setting, an interaction of DSG2 and EGFR has been observed (Ungewiss et al., 2018). However, to our knowledge in cardiomyocytes this complex was not known before. It has been very well demonstrated that desmosomes in the epithelia or ICDs in the cardiomyocytes not only provide cellular cohesion but can also be part of a signaling hub (Broussard et al., 2015, Najor et al., 2017, Muller et al., 2021, Yeruva and Waschke, 2022), thus disrupting EGFR signaling in this context might lead to changes at the desmosome.

On a single molecule level in an AFM setting, we found that binding events mediated by DSG2, but not DSC2 and N-CAD were increased after erlotinib treatment in HL-1 cardiomyocytes. Increased binding frequency at the cell borders indicates an increased binding partner localization towards the cell borders. However, it might also be that active EGFR is disturbing the interaction possibility of the binding partner with the DSG2-coated AFM tip, which is restored upon EGFR inhibition. In a cell-free AFM setting DSG2 does not only interact homophilically with DSG2, but also heterophilically with DSC2 (Fuchs et al., 2022). In the AFM setting, only interactions with “free” proteins can be measured. Since we observed an increase of DSG2 at the cell borders upon EGFR inhibition, we suppose that the increased binding frequency at cell borders, in line with our results from the Ca^2+^-switch assays indicate increased desmosomal assembly, where freshly assembled desmosomal precursors are available for interaction with the opposite cell DSG2 or DSC2. At the same time, the decrease of DSC2 binding frequency might indicate that either DSC2 or its binding partners are not available for interactions with the DSC2-coated tips, especially at the cell surface. Since we did not observe any significant changes in the DSC2 binding frequency at the cell border, we assumed that the observed decrease in DSC2 binding frequency at cell surface was not off significant concern.

When we knocked down *Egfr* by siRNA, SRC inhibition was still effective in inducing positive adhesiotropy, indicating that the SRC inhibition-mediated effect on cardiomyocyte cohesion is either downstream or at least in part independent of EGFR. However, Western blot analyses revealed that PP2 also caused a significant decrease in EGFR phosphorylation at Y845 and Y1068, indicating SRC acts upstream of EGFR. We have previously shown in keratinocytes that SRC via its target cytoskeletal protein cortactin contributes to the loss of cell cohesion mediated by DSG3 autoantibodies (Kugelmann et al., 2019). Similarly, in cardiomyocytes SRC might also have a direct interaction with desmosomal proteins, independent of the EGFR molecule. Infact, in intestinal epithelial cells DSG2, EGFR and SRC were found to be in one complex (Ungewiss et al., 2018). Upon knockdown of *DSG2*, DSC2 has been shown to be upregulated in colonic adenocarcinoma cell lines (Kamekura et al., 2014), which might explain why we can still observe a positive adhesiotropic effect of EGFR or SRC inhibition upon *Dsg2* knockdown. On the other hand, the remaining DSG2 upon *Dsg2* knockdown could be sufficient to induce positive adhesiotropy after EGFR inhibition. Indeed, we never observed a decrease in cell cohesion upon *Dsg2* knockdown (Schinner et al., 2020, Shoykhet et al., 2020) which further confirms that the partial presence of DSG2 can provide mechanical stability to the cells.

Further, we observed an increased recruitment of DSG2 and DP to the membranes of HL-1 cardiomyocytes, increasing area composita length. The longer areae compositae in HL-1 cardiomyocytes could be the cellular correlate of increased thickness of DSG2 and DP staining in murine cardiac slices. A similar reorganization of ICDs has already been observed in electron microscopy upon adrenergic signaling, which also induces positive adhesiotropy in cardiomyocytes (Yeruva et al., 2020). We found that the recruitment of DSG2 and DP into the cellular junctions is not caused by increased mobility, rather by an enhanced desmosomal assembly, which is supported by increased DES insertions into DP. Interestingly, in *Jup*^-/-^ mice, we observed changes in DSG2 but not DP localization at the ICD, indicating that the effect of EGFR and SRC inhibition in intact cardiac tissue is at least in part mediated by PG. Indeed, it was shown previously that in *Jup*^-/-^ keratinocytes, steady-state protein levels and recruitment of desmosome components to the membrane were reduced (Yin et al., 2005).

Mechanistically, PG-phosphorylation at tyrosine residues was shown to be involved in EGFR-induced loss of cell adhesion, which decreased association of PG to DP (Gaudry et al., 2001), and triggered translocation of PG and DP from the membrane towards the cytoplasm (Gaudry et al., 2001, Bektas et al., 2013). Furthermore, EGF-induced phosphorylation of PG could be prevented by EGFR inhibition (Lorch et al., 2004). In consensus with the above studies, an increase in co-immunoprecipitation of PG and PKP2 proteins with DP in HL-1 cardiomyocytes after inhibition of EGFR or SRC was observed in this study. These results further strengthen our hypothesis that desmosomal assembly is increased upon EGFR or SRC inhibition.

Similar to our findings in this study, in keratinocytes, inhibition of ROCK increased cellular migration and decreased cellular cohesion (Jackson et al., 2011, Rotzer et al., 2014). In corneal epithelial cells, inhibition of ROCK led to a translocation of E-Cadherin to the cytoplasm (Anderson et al., 2002). Apart from that, using an *in silico* mathematical model, it has been suggested that ROCK plays a role in the pathogenesis of AC (Parrotta et al., 2021). Furthermore, inhibition of ROCK during early cardiac development did lead to AC in a murine model with a reduced localization of PG at the ICD (Ellawindy et al., 2015). An involvement of RhoA, which is upstream of ROCK, in functional desmosomal assembly has been described in squamous carcinoma cell lines (Godsel et al., 2010). Here, we observed that ROCK inhibition decreased cellular cohesion, but did not alter DSG2 or DP localization to the cell borders significantly in HL-1 cardiomyocytes, though a decreasing nonsignificant trend could be observed. Nonetheless, ROCK inhibition decreased erlotinib induced DP and DSG2 localization to the cell borders. Furthermore, after a Ca^2+^-switch, ROCK inhibition alone did not alter the cellular cohesion, whereas it prevented the enhanced desmosomal assembly induced by erlotinib. We therefore believe that erlotinib mediated desmosomal assembly and cohesion in cardiomyocytes requires ROCK. We suppose that a possible effect of ROCK inhibition on cellular cohesion after a Ca^2+^-switch was below the detection limit of dissociation assays. These results warranty further studies to understand the role of ROCK in the pathomechanisms of AC.

Based on our results, we anticipate that short-term EGFR inhibition can be beneficial in the setting of AC. Furthermore, unlike other EGFR inhibitors, reports of adverse cardiac effects after erlotinib treatment are very rare (Kenigsberg et al., 2017, Stuhlmiller et al., 2017, Barish et al., 2019, Nagashio et al., 2021). Taken together, we conclude that inhibiting EGFR, thereby enhancing desmosome assembly via ROCK stabilizes desmosome integrity and cardiomyocyte cohesion. However, further studies, investigating the *in vivo* effect of EGFR inhibition utilizing AC mouse models as well as the effects on electromechanical coupling are needed, which are beyond the scope of current study. Nevertheless, we believe that our study is a first step that paves the way for future studies targeting EGFR inhibition or ROCK activation by erlotinib as a treatment option for AC.

## Materials and Methods

### Cell lines

HL-1 cells, which are immortalized female murine atrial myocyte cells, provided by William Claycomb (LSU Health Sciences Center, New Orleans, USA), were cultured in supplemented medium as described before (Shoykhet et al., 2020). Cells were cultured in medium containing norepinephrine, while for experiments, medium without norepinephrine was used.

### Murine cardiac slice culture

Animal handling was in accordance with the guidelines from the Directive 2010/63/EU of the European Parliament and approved by the regional government of Upper Bavaria (Gz. ROB-55.2-2532.Vet_02-19-172). Previously characterized 12 week old C57BL6/J wildtype and cardiomyocyte specific *Jup*^-/-^ mice were used for experiments (Schinner et al., 2017, Schinner et al., 2020). In brief, mice with loxP sites flanking exon 1 of the junctional plakoglobin gene (*Jup*) on both alleles (Juptm1.1Glr/J mice, the Jackson Laboratory, Bar Harbor, USA) were bred with mice heterozygous for loxP sites in *Jup* with an expression of the recombinase Cre under the control of cardiac-specific alpha myosin heavy chain promoter (Myh6), which were generated by crossing with B6.FVB-Tg(Myh6-cre)2182Mds/J mice (The Jackson Laboratory).

Murine cardiac slice cultures were obtained as described before (Shoykhet et al., 2020) with minor adjustments. In brief, after anesthetizing mice with isoflurane, mice were sacrificed by cervical dislocation, the hearts were collected, put into pre-cooled slicing buffer, embedded in low-melt agarose and sliced with a vibratome (LeicaVT1200S vibratome) into 200 µm sections. For treatments, consecutive slices were used to minimize slice-to-slice variability. After respective treatments, slices were processed for lysate preparation, dispase-based dissociation assays and immunostainings. For immunostainings, slices were embedded in NEG-50™ frozen section medium and stored at -80°C until further processing.

### Mediators

Samples were treated with mediators for 90 minutes. Erlotinib (Santa Cruz, #sc-202154) was used as an EGFR inhibitor at a final concentration of 2.5 µM. PP2 (Merck, #529573) was used to inhibit the EGFR signaling pathway by inhibiting SRC at a final concentration of 10 µM. Y27632 was used to inhibit ROCK at a final concentration of 10 µM (Tocris, #1254). When a Ca^2+^-switch was performed, cells were treated with 5 mM EGTA (VWR, #0732) for 90 minutes. Then, medium was changed back to Ca^2+^-containing medium for 90 minutes and mediators were added at the same time.

### Lysate preparation

For Western blot analyses, lysates were prepared in SDS lysis buffer with protease and phosphatase inhibitors (cOmplete™ Protease Inhibitor Cocktail, Roche, #CO-RO and PhosStop™, Roche, #PHOSS-RO). In the case of HL-1 cardiomyocytes, the medium was removed after treatments and cells were kept on ice, washed with PBS, scraped into SDS lysis buffer and sonicated. When working with murine cardiac slice cultures, slices were treated, washed with TBS and snap-frozen in liquid nitrogen. To prepare the tissue lysates, SDS lysis buffer was added, the samples were transferred into gentleMACS™ M-tubes (Miltenyi Biotec, #130-093-236) and dissociated using the protein_01_01 program of the gentleMACS™ OctoDissociator (Miltenyi Biotec, #130-095-937).

For co-immunoprecipitation experiments, cells were cultured in T75 flasks. After treatments, RIPA lysis buffer (10 mM Na_2_HPO_4_, 150 mM NaCl, 1% Triton X-100, 0.25% SDS, 1% sodium deoxycholate, pH=7.2) containing protease and phosphatase inhibitors was added to the cells and incubated for 30 minutes on ice on a rocking platform. The lysate was then transferred to gentleMACS™ M-tubes, and dissociated using the protein_01_01 program of the gentleMACS™ OctoDissociator.

For Triton assays, cells were washed with ice cold PBS and cells were kept on ice on a rocking platform in Triton buffer (0.5% Triton X-100, 50 mM MES, 25 mM EGTA, 5 mM MgCl_2_, pH=6.8) with protease and phosphatase inhibitors for 20 minutes. After that, the lysate was transferred to Eppendorf tubes and centrifuged at 4°C for 10 minutes at 15000 g in an Eppendorf 5430R centrifuge. The supernatant containing the Triton-soluble proteins was transferred to a new reaction tube, while the pellets were washed with Triton buffer, dissolved in SDS lysis buffer and sonicated.

For the PamGene Kinase assay, after the respective treatments cells were washed twice with ice-cold PBS and lysed into M-PER™ lysis buffer (Thermo Fisher #78503) supplemented with Halt™ Protease and Phosphatase inhibitors (Thermo Fisher #87786 and #78420). The lysate was transferred to Eppendorf tubes, centrifuged at 4°C for 15 minutes at 15000 g in an Eppendorf 5430R centrifuge and the supernatant was aliquoted into fresh tubes, snap-frozen in liquid nitrogen and stored at -80°C until further analysis.

For all assays, protein concentration was measured using the Pierce™ Protein Assay Kit (Thermo Fisher, #23225) according to the manufacturer’s protocol.

### Western blot analyses

Lysates were denatured in Laemmli buffer and boiled for 10 minutes at 95°C, before being loaded on SDS-PAGE gels. The PageRuler™ Plus Prestained Protein Ladder (Thermo Fisher, #26620) was used as a marker. After electrophoresis, proteins were transferred to a nitrocellulose membrane (Thermo Fisher, #LC2006) using the wet-blot method. Membranes were then blocked in 5% milk/TBST, in case of anti-phospho-EGFR antibodies in 5% BSA/TBST, or in case of anti-DP antibodies in ROTI®Block (Carl Roth, #A151.2). For Western blots with lysates from murine tissue samples, as well as Triton blots, the No-Stain™ Protein Labeling Reagent (Thermo Fisher, #A44449) was used as loading control according to the manufacturer’s protocol. Following primary antibodies were added overnight at 4°C on a rocking platform: anti-DP (Progen, #61003), anti-DSG1/2 (Progen #61002), anti-phospho-EGFR Y845 (Cell Signaling #2231), anti-phospho-EGFR Y1068 (Cell Signaling, #2234), anti-EGFR (sc-373746 (for HL-1 cardiomyocytes) and ab52894 (for murine tissue)), anti-phospho-ERK1/2 ((E-4) Santa Cruz, #sc-7383), anti-ERK (Cell Signaling, #9102), anti-phospho-PG (S665, self-made as described before (Schinner et al., 2020)), anti-PG ((PG5.1) Progen #61005), anti-PKP2 (Progen, #651167) and anti-α-tubulin ((DM1A), abcam, #ab7291). After washing, membranes were incubated with species-matched, HRP-coupled secondary antibodies (Dianova, #111-035-045 or #115-035-068) diluted in TBST or 5% milk/TBST in case of phospho-EGFR antibodies, and developed using an iBright™ FL1500 (Thermo Fisher) developer with SuperSignal™ West Pico PLUS Chemiluminescent Substrate (Thermo Fisher, #34577). All primary antibodies were used at 1:1000 dilutions, apart from phospho-EGFR and EGFR antibodies (1:500), phospho-PG (1:20), anti-PKP2 (1:25) and anti-α-tubulin (1:4000). Quantification was performed using the Image Studio Lite v.5.2 or the iBright™ Analysis Software (Invitrogen).

### Dissociation Assays

Dissociation assays in HL-1 cardiomyocytes were performed as described before (Shoykhet et al., 2020). In brief: after treatment, medium was removed and Liberase-DH (Sigma-Aldrich, #LIBDH-RO) was added, followed by dispase II (Sigma-Aldrich, #D4693). Cells were incubated at 37°C, 5% CO_2,_ until the cell monolayers detached from the well bottoms. Then, the enzyme mix was carefully replaced by HBSS and mechanical stress was applied by shaking at 1.31 g for 5 minutes on an orbital shaker (Stuart SSM5 orbital shaker). For better visibility, MTT (Sigma-Aldrich, #M5655) was added to the cells and images were taken to count the number of fragments. The amount of fragmentation serves as an indirect measurement of intercellular cohesion, as fewer fragments suggest a stronger cellular cohesion.

Dissociation assays in murine cardiac slices were performed similarly to dissociation assays in HL-1 cardiomyocytes. However, in this case, Liberase-DH and dispase II were added simultaneously for 30 minutes. MTT was added and equal mechanical stress was applied to all samples of an experiment using an electrical pipette. Then, the content of each well was filtered using a 70 µm nylon membrane (PluriSelect, #43-10070-60), and transferred to a gridded 96-well plate. Pictures of the well were taken and stitched together using the AutoStich software (Brown, 2018). For quantification, only the number of viable dissociated cardiomyocytes was counted, distinguishable by their rod shape and the MTT staining. To control variations due to different slice size or location in the ventricle, consecutive slices for control and treatments were used, and treatments were normalized to the respective control slice.

### Immunostainings

Cells were seeded on coverslips, and fixed with paraformaldehyde (PFA) after respective treatments. 7 µm thick sections of murine cardiac slices were cut with a cryostat (HM500OMV, Microm, Walldorf, Germany), sections were transferred to glass slides and adhered by warming up to 38°C. After PFA fixation, the samples were permeabilized with 0.1% Triton-X 100 and blocked with bovine serum albumin and normal goat serum (BSA/NGS). Primary antibodies diluted in PBS were added and incubated overnight at 4°C in a wet chamber. Following primary antibodies were used: anti-DES (abcam, #ab32362), anti-DP (Progen, #61003), anti-DSG2 (Progen, #610121) and anti-N-CAD (BD Transduction Laboratories™, #610921). When working with HL-1 cardiomyocytes, PBS was used for washing steps, while for murine tissue samples, 50 mM NH_4_Cl in PBS was used. After washing, species-matched, fluorophore-coupled secondary antibodies ((anti-mouse Alexa Fluor® 488 (Dianova, #115-545-003), anti-mouse Cy3 (Dianova, #115-165-164) or anti-rabbit Cy5 (Dianova, #111-175-144)) were added for 60 minutes. In the case of wheat germ agglutinin (WGA) or Phalloidin staining (Thermo Fisher, #W11261 and Thermo Fisher, #A12379), the dye was added together with the secondary antibodies. WGA was used at 2 µg/ml and Phalloidin at 130 nM concentrations. During the last 10 minutes of secondary antibody incubation, DAPI (Sigma Aldrich, #D9542, 0.5 µg/ml) was added. After washing, coverslips were mounted on microscope slides using NPG (Sigma-Aldrich, #P3130). Slides were analyzed using a Leica SP5 II confocal microscope (Leica, Mannheim, Germany) with a 63X oil objective with a numerical aperture of 1.4 using Immersol 518F (Zeiss). Images were acquired at room temperature with the LAS-AF software. Z-scans were performed at 0.25 µm thickness spanning the whole cell volume. Colocalization analyses were performed as described before (Shoykhet et al., 2020), in brief: the membrane region was marked as a region of interest (Supplementary Figure 5A). In this region, the amount of stained pixels in the other channel, where colocalization was to be quantified, was measured and a ratio of colocalization at the membrane was calculated. Measurement of staining width in immunostainings performed in murine cardiac slice cultures, is exemplified in Supplementary Figure 5B.

For stimulated emission depletion (STED) immunostainings, a slightly modified staining protocol was used: instead of washing with PBS, cells were washed with 50 mM NH_4_Cl in PBS. Primary antibodies were diluted in BSA/NGS and the following secondary antibodies were used: STAR RED anti-rabbit (abberior, #STRED-1002) and Alexa Fluor® 594 anti-mouse (abcam, #ab150120), both 1:200 dilution. Slides were mounted with ProLong™ Diamond Antifade (Thermo Scientific, #P36961). Samples were analyzed at room temperature using a STED expert line microscope from Abberior with a 100X oil objective with a numerical aperture of 1.4 using IMMOIL-F30CC (Olympus). Images were acquired with a DMK33G274 camera and the Imspector software. For quantification, areas with a “railroad-like” staining of DP were considered as area composita. To quantify DES insertions into the area composita, the number of spots where DES filaments were inserting into the area composita were counted.

### Immunoprecipitation

For pulldowns, 1 µg of respective antibodies were added to 1.5 mg of protein lysate overnight, and kept at 4°C on a spinning wheel. To rule out unspecific antibody or bead binding, IgG control samples were prepared, in parallel. Therefore, the same amount of protein lysate was used as for pulldowns, however, instead of primary antibody, 1 µg of normal mouse IgG (Merck, #12-371) was added to the samples. Pulldowns were performed with Protein G Dynabeads™ (Thermo Fisher, #10004D), by adding the lysate-antibody mix for 1 hour at 4°C on a spinning wheel to beads prewashed in RIPA buffer with protease and phosphatase inhibitors. Washing steps were performed using a magnetic rack. Then, beads were washed twice with wash buffer 1 (50 mM Tris-HCl, 150 mM NaCl, 0.1 mM EDTA, 0.5% Tween20, pH=7.5), thrice with wash buffer 2 (100 mM Tris-HCl, 200 mM NaCl, 2 M Urea, 0.5% Tween20, pH=7.5), and twice with 1% Triton X-100 in PBS. All wash buffers were supplemented with protease and phosphatase inhibitors. After the last wash step, beads were resuspended in Laemmli buffer, denatured for 10 minutes at 95°C, the lysate was separated from the beads using a magnet and loaded on a SDS-PAGE gel for Western blot analyses. 20 µg of total lysate was used as input control for the Western blots.

### Atomic Force Microscopy

Atomic Force Microscopy (AFM) measurements were performed as described before (Schinner et al., 2017, Schinner et al., 2020) using a Nanowizard® III AFM (JPK instruments, Berlin, Germany) with an optical fluorescence microscope (Axio Observer D1, Carl Zeiss) at 37°C. Cantilevers (MLCT AFM Probes, Bruker, Calle Tecate, CA) were functionalized with recombinant proteins as described elsewhere (Ebner et al., 2007). The following recombinant proteins were used: DSC2-Fc, DSG2-Fc (both self-made by Protein A purification of cell culture supernatants from overexpressing CHO cells, as described in (Waschke et al., 2005)) or recombinant N-CAD-Fc (R&D Systems, #6626-NC-50). The canonical pyramidal-shaped D-tip was used. The thermal noise method (Hutter and Bechhoefer, 1993) was used to determine the spring constants of the tips in order to obtain correct values for unbinding forces. For experiments, HL-1 cardiomyocytes were seeded on 24 mm glass coverslips. A bright field image of the cells of interest was acquired with a 63X objective. Afterwards an overview picture of an area of interest was taken in the force-based quantitative imaging (QI) mode with the following settings: pulling speed: 50 µm/s, setpoint: 0.2 nN, z-length 2.5 µm. Based on the QI image, areas for adhesion measurements were chosen (Supplementary Figure 5C). Adhesion measurements were performed on cell borders as well as cell surfaces on an area of 1.5 µm x 5.0 µm using the force mapping mode with the following settings: relative setpoint: 0.2 nN, z-length: 2.5 µm, extension speed: 5.0 µm/s, extend and retract delay: 0.1 s. Information of approach and retraction forces was recorded in force-distance curves (FDCs). The resulting FDCs were analyzed using the JPKSPM Data processing software. Binding events are visible by a step in the FDCs, representing the force needed to rupture the binding between the functionalized tip and the binding partner on the cell. Hence, this force is the unbinding force. Unbinding forces were analyzed using the extreme fit distribution in Origin 2017 software.

### siRNA-mediated knockdown

For siRNA-mediated knockdown, cells were transfected with a transfection reagent, comprised of OptiMEM (Thermo Scientific, #31985070), RNAiMAX (Thermo Scientific, #13778150) and the ON-Target siRNA (Dharmacon, siNT #D-001810-10, si*Egfr* L-040411-00-0005, si*Dsg2* #L-042514-01-005, si*Dsp* #L-040653-01-005) according to manufacturer’s protocol. Cells were transfected for 24 hours. 72 hours post transfection, medium was changed again. Experiments were performed 84 hours post-transfection. Each knockdown was confirmed by Western blot.

### Fluorescence Recovery After Photobleach

For Fluorescence Recovery After Photobleach (FRAP) measurements, HL-1 cardiomyocytes were seeded in ibidi 15 µ-Slide 8 Well Glass Bottom chambers (ibidi, #80827). 24 hours after seeding, cells were transfected with 1 µg DSG2-GFP construct as described before (Schinner et al., 2017), using TurboFect (Thermo Fisher, #R0531). Images were captured with a Leica SP5 confocal microscope (Leica, Mannheim, Germany) with the LAS-AF software starting 60 minutes after treatment with the vehicle or erlotinib. Using the Leica FRAP Wizard, 5 pictures were taken before bleaching, then an area of interest was bleached with 20 pulses at 100% laser intensity for 0.523 s. After that, 60 pictures were taken every second, 20 pictures were taken every 3 seconds and 10 pictures were taken every 10 seconds. Halftime of recovery, as well as immobile fractions, were calculated using the Leica FRAP Wizard.

### Quantification and statistical analysis

Images were processed with Adobe Photoshop CS5, Image Studio Lite v.5.2 and ImageJ software. The cartoon was created using BioRender. Statistical analyses were performed using GraphPad Prism 8. Student’s *t*-tests or 1-Way ANOVA with Post-hoc tests were applied after outlier removal and are explained in figure legends. Data are represented as mean ± standard deviation. For dissociation assays as well as for quantification of Western blots in HL-1 cardiomyocytes, values were normalized to the average control value of all experiments. For dissociation assays in murine cardiac slices, the number of dissociated cells was normalized to the respective control slice of the same mouse. Significance was assumed for p ≤ 0.05.

### PamGene Kinase Assay

The Serine-Threonine Kinase (STK) and Protein Tyrosine Kinase (PTK) microarray assays were performed according to the manufacturer’s instructions (Protein Tyrosine Kinase assay on PamStation^®^ 12 User Manual, Version 3.0; Serine Threonine Kinase assay on PamStation^®^ 12 User Manual, Version 5.1) and as described earlier (Zillikens et al., 2021). In brief, cell lysates were combined with reaction mixtures produced with the contents of the STK reagent kit (PamGene, Cat. No. 32201) or PTK reagent kit (PamGene, Cat. No.32112) and ampuwa ultrapure water (Fresenius Kabi) and placed on Serine threonine Kinase PamChips (PamGene, Cat. No. 32501) and Protein Tyrosine Kinase PamChips (PamGene, Cat. No. 32508) respectively. Cell lysates originating from the same culture passage and experimental setup (n=3) were processed within the same assay run to enable accurate comparisons. FITC-conjugated anti-phosphotyrosine and anti-phosphoserine antibodies were used for visualization during and after the pumping of lysates through the three-dimensional surface of the array. The capture of substrate phosphorylation signals was enabled by a computer-controlled CCD camera and measured repeatedly during a 1-hour kinetic protocol using the Evolve software (PamGene International B.V.). The analysis of the images was performed using the BioNavigator Software (Ver. 6.3), with the predesigned protocols “STK Image Analysis“, “PTK Image Analysis“, “STK Basic Processing“ and “PTK Basic Processing“ and “Log Fold change (No Log). Additional applications used were “Fit and apply a combat model“, “STK upstream kinase analysis“ and “PTK upstream kinase analysis“. After visual check and quality control, endpoint signal intensities minus background signals were calculated by BioNavigator for each spot representing each kinase peptide substrate per array.

Subsequently, the data were log_2_-transformed before mean replicate signal intensity within each experiment was calculated for each peptide substrate. A kinase was considered to be modulated (either activated or inhibited) if it had a mean specificity score (negative decadic logarithm of the likelihood of obtaining a higher difference between the groups when assigning peptides to kinases randomly) of 1 and a significance score (likelihood of obtaining a higher difference for random assignment of values to treatment- and control groups ) of 0.5. To detect changes caused by Erlotinib, samples for each timepoint were compared with the vehicle control samples for the same timepoint. The mean kinase statistic (calculated by averaging the difference between the signal intensity of a sample and its control value, normalized against a pooled estimate of the standard deviation in each sample, for each peptide assigned to a specific kinase) was used for further analysis.

## Online Supplementary Material

Figure S1 shows Western blot confirmations of the erlotinib and PP2 effects on HL-1 cardiomyocytes. Figure S2 shows representative images of the dispase-based dissociation assay in HL-1 cells after erlotinib or PP2 treatments. Figure S3 shows an increased DES insertion into DP upon erlotinib or PP2 treatments. Figure S4 shows that EGFR or SRC inhibition do not cause changes in protein localization between detergent soluble and insoluble fractions. Figure S5 shows exemplified immunostaining quantifications and the area choice in the AFM measurements.

Table S1 shows the results of the PamGene assay.

## Author contributions

MS, OD, PM, MH and CO acquired the data. MS and SY analyzed the data and drafted the manuscript. All authors proof-read the manuscript. RJL supervised the PamGene assay, JW, SY handled supervision of the project. JW and SY made critical revision of the manuscript for important intellectual content and designed the research.

## Acknowledgements

This work was supported by the Ludwig-Maximilian-University Munich through the FöFoLe program, the Deutsche Forschungsgemeinschaft (grant WA2474/11-1 to JW, EXC 2167, GRK 2633, and LU 877/23-1 to RJL) as well as by the Schleswig-Holstein Excellence-Chair Program from the State of Schleswig Holstein, and grant CRSII5_202301/1 from the Swiss National Science Foundation. The authors declare no competing financial interests. We thank Martina Hitzenbichler, Daniela Kugelmann, Kathleen Plietz, Kilian Skowranek and the AFM tip coating team for their help and technical assistance.

## Abbreviation

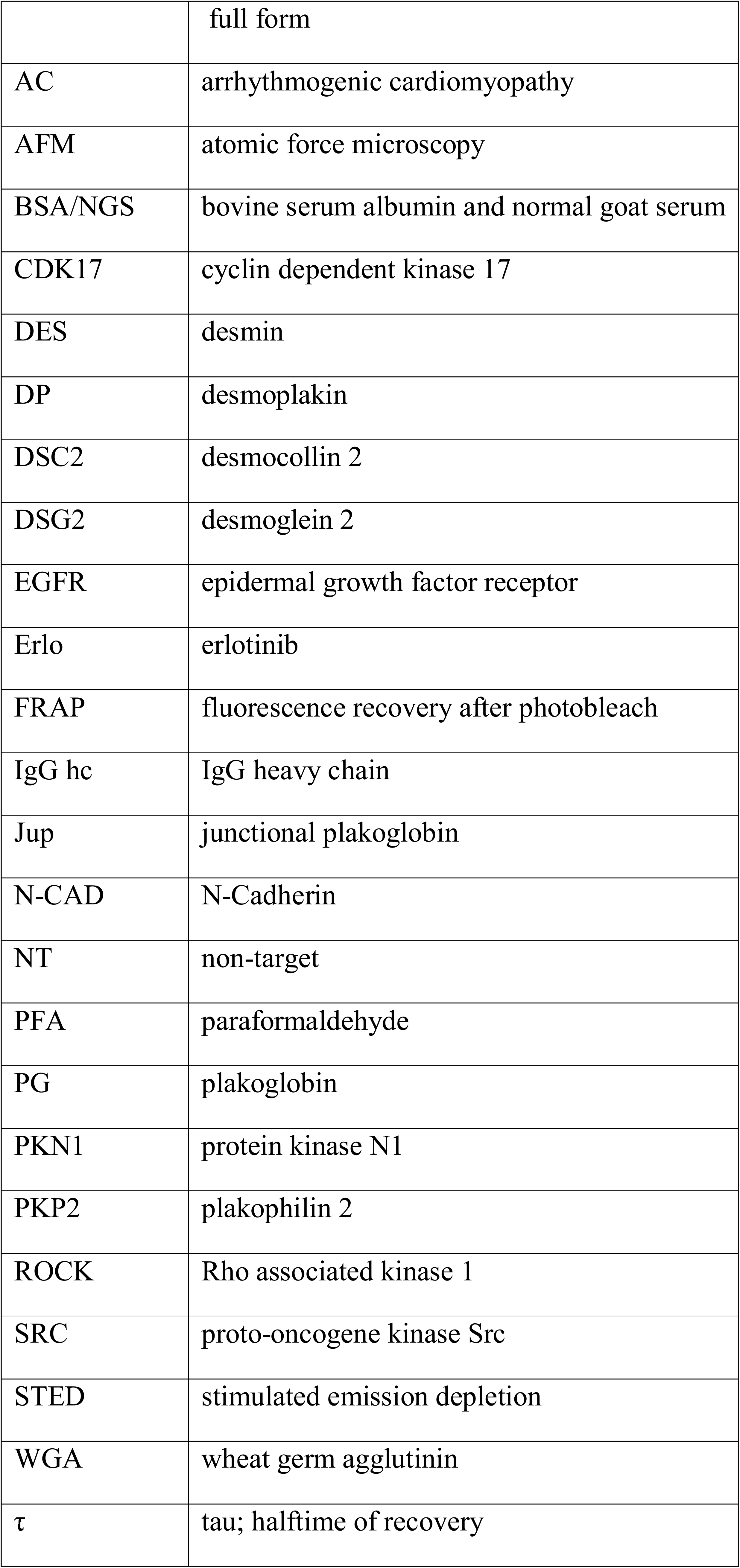

