## Supplementary data for "EGFR inhibition led ROCK activation enhances desmosome assembly and cohesion in cardiomyocytes"

Figure S1 – confirmation of erlotinib and PP2 effect

**A:** Quantification of Western blots showing protein expression of EGFR in murine cardiac slices from *Jup*^+/+^ and *Jup*^-/-^ mice, * p ≤ 0.05 Student’s *t*-test (p=0.007), N=6 mice**. B-D**: Representative Western blots showing protein expression of phospho-EGFR (**B**: Y845 and **C**: Y1068) and EGFR and **D**: phospho-ERK1/2 and ERK in HL-1 cardiomyocytes after treatments with erlotinib or PP2. * p ≤ 0.05 (pEGFR845 p=0.4916 for DMSO vs erlotinib and p=0.0154 for PP2; pEGFR1068 p=0.9743 for DMSO vs erlotinib and p=0.0495 for DMSO vs PP2, pERK1 p=0.0196 for DMSO vs erlotinib and p=0.0138 for DMSO vs PP2, pERK2 p=0.5008 for DMSO vs erlotinib and p=0.5321 for DMSO vs PP2), 1-way ANOVA with Holm-Sidak correction, N=5 biological replicates.

Figure S2 – Positive adhesiotropy induced by EGFR or SRC inhibition is dependent on DP

Representative pictures of the wells of the dispase-based dissociation assay in HL-1 cardiomyocytes after siRNA-mediated knockdown of **A**: EGFR, **B**: DSG2 and **C**: DP.

Figure S3 – Increased DES insertion into DP upon erlotinib or PP2 treatments

Representative STED images of HL-1 cardiomyocytes stained for DP and DES. Bar graphs represent percentage of desmosomes with Desmin insertions. Scale bar in overviews: 10 µm, scale bar in zooms: 2 µm. *p ≤ 0.05 (p=0.0147 for DMSO vs erlotinib and PP2), 1-way ANOVA with Holm-Sidak correction, N=4 biological replicates.

Figure S4 – EGFR or SRC inhibition do not cause changes in protein localization between detergent soluble and insoluble fractions

**A**: Representative Triton assay blots for DP, EGFR, PG and DSG2, NoStain™ dye served as a loading control. Triton soluble fraction represents cytosolic proteins and membrane-bound proteins in Triton insoluble fraction. **B**: Quantifications of Triton assay blots for DP, EGFR, PG and DSG2 * p ≤ 0.05 (DP insoluble: p=0.5667 for DMSO vs erlotinib and p=0.9298 for DMSO vs PP2; DP soluble: p=0.8449 for DMSO vs erlotinib and DMSO vs PP2; EGFR insoluble: p=0.6873 for DMSO vs erlotinib and p=0.6431 for DMSO vs PP2; EGFR soluble: p=0.9298 for DMSO vs erlotinib and p=0.644 for DMSO vs PP2; PG insoluble: p=0.8289 for DMSO vs erlotinib and p=0.1936 for DMSO vs PP2; PG soluble: p=0.7772 for DMSO vs erlotinib and p=0.159 for DMSO vs PP2; DSG2 insoluble: p=0.7072 for DMSO vs erlotinib and p=0.4255 for DMSO vs PP2; DSG2 soluble: p=0.578 for DMSO vs erlotinib and p=0.3486 for DMSO vs PP2) 1-way ANOVA with Holm-Sidak correction, N=5-7 biological replicates. **C**: Representative pictures of the wells of the dispase-based dissociation assay in HL-1 cardiomyocytes after 90 minutes of Ca^2+^-depletion and subsequent Ca^2+^-repletion together with erlotinib PP2.

Figure S5 – Methods and quantifications explained

**A**: Quantification method for colocalization in immunostainings. The region of interest was chosen using N-CAD, and transferred to the image of the DSG2 staining. In this region, the amount of stained pixels was measured and a ratio DSG2 to N-CAD at the membrane was calculated. **B**: Quantification method for DP or DSG2 immunostaining width in murine cardiac slices. The width of respective protein staining was measured in two areas per ICD, the average values were calculated. WGA staining was used in order to orient the ICDs. Staining width was measured in areas, where WGA, DP and DSG2 stainings colocalized. The measurements were performed using ImageJ as shown. **C**: Representative QI topography image of HL-1 cells with chosen areas for AFM measurements at cell borders and cell surfaces.

Table S1

PamGene assay after 15, 30 and 60 minutes of erlotinib treatment. Samples were compared to respective control-treated specimen. N=3 biological replicates. Kinase activity was considered to be altered, when the mean specificity score was above 1 and the significance score above 0.5.
