## Supplementary figures and images for "EGFR inhibition led ROCK activation enhances desmosome assembly and cohesion in cardiomyocytes"

### Supplementary figure 1

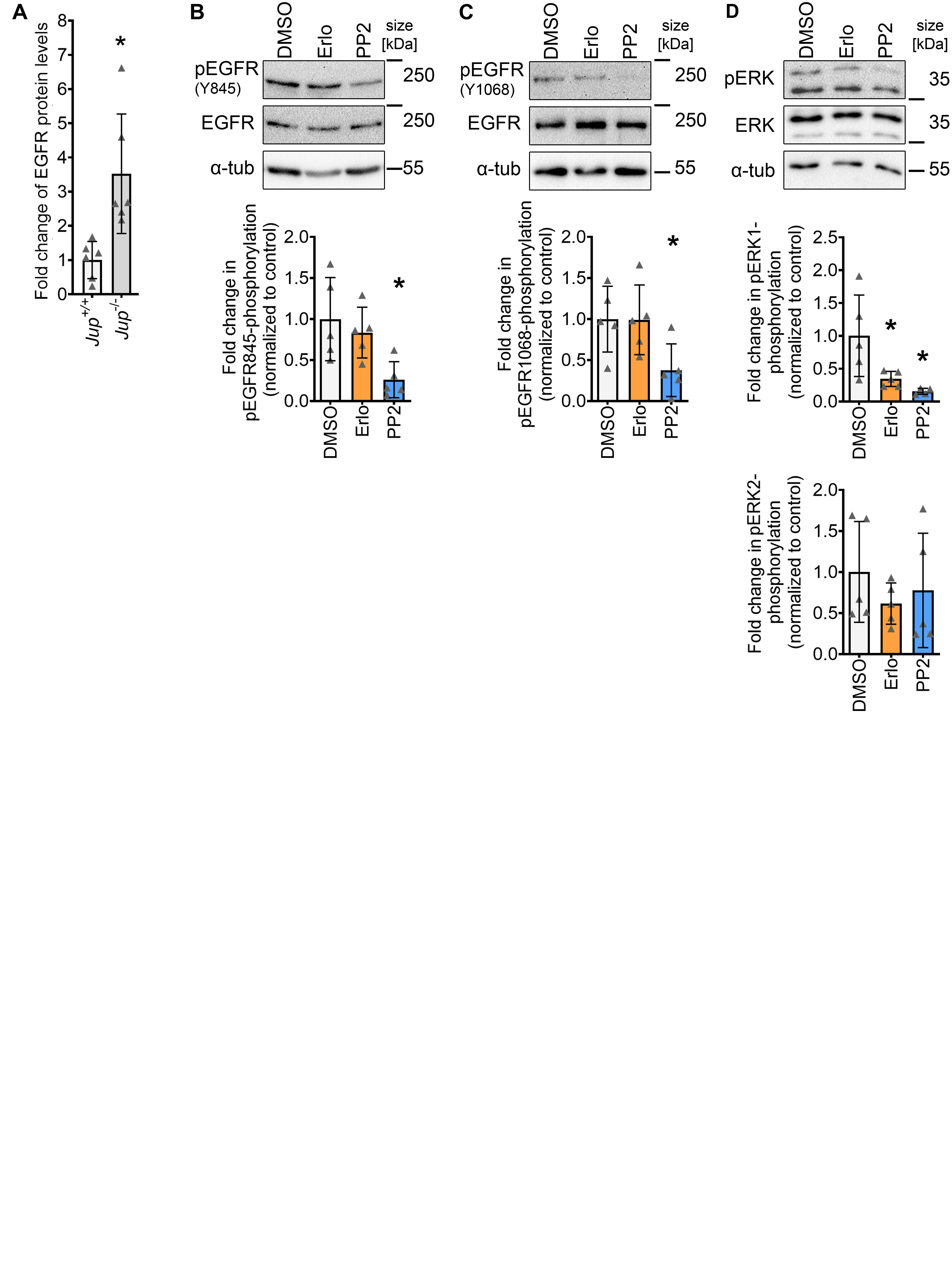

### Supplementary figure 2

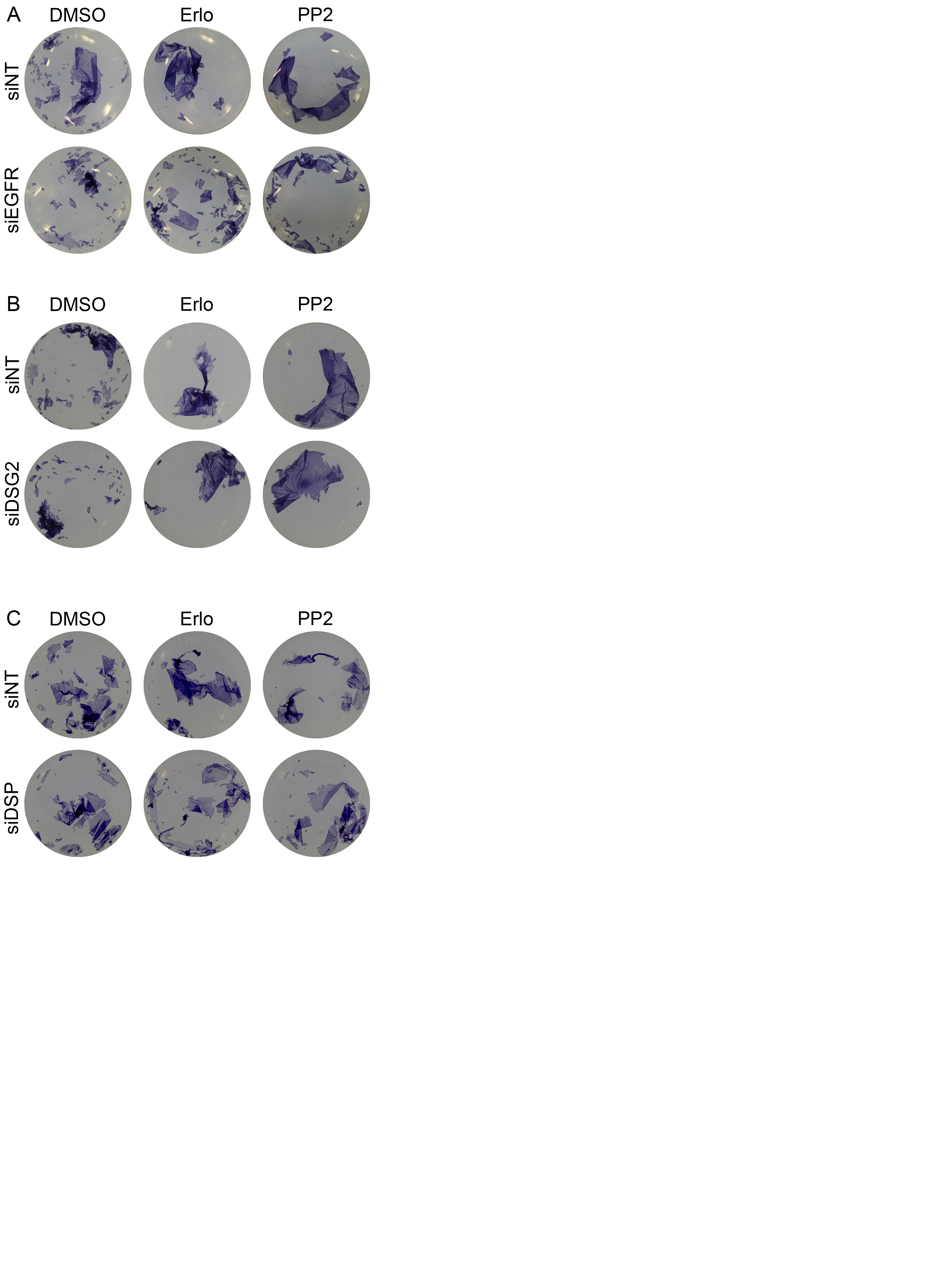

### Supplementary figure 3

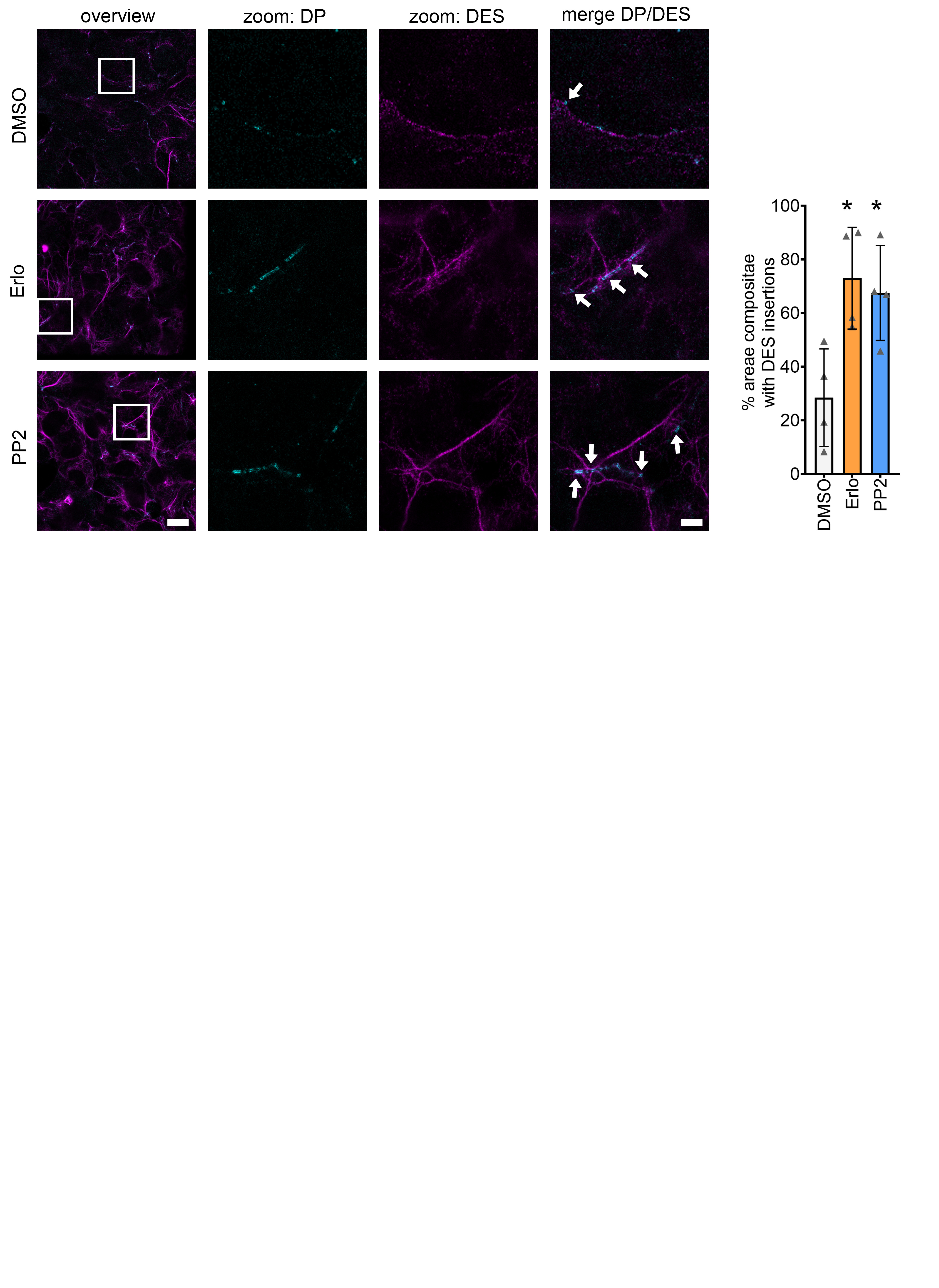

### Supplementary figure 4

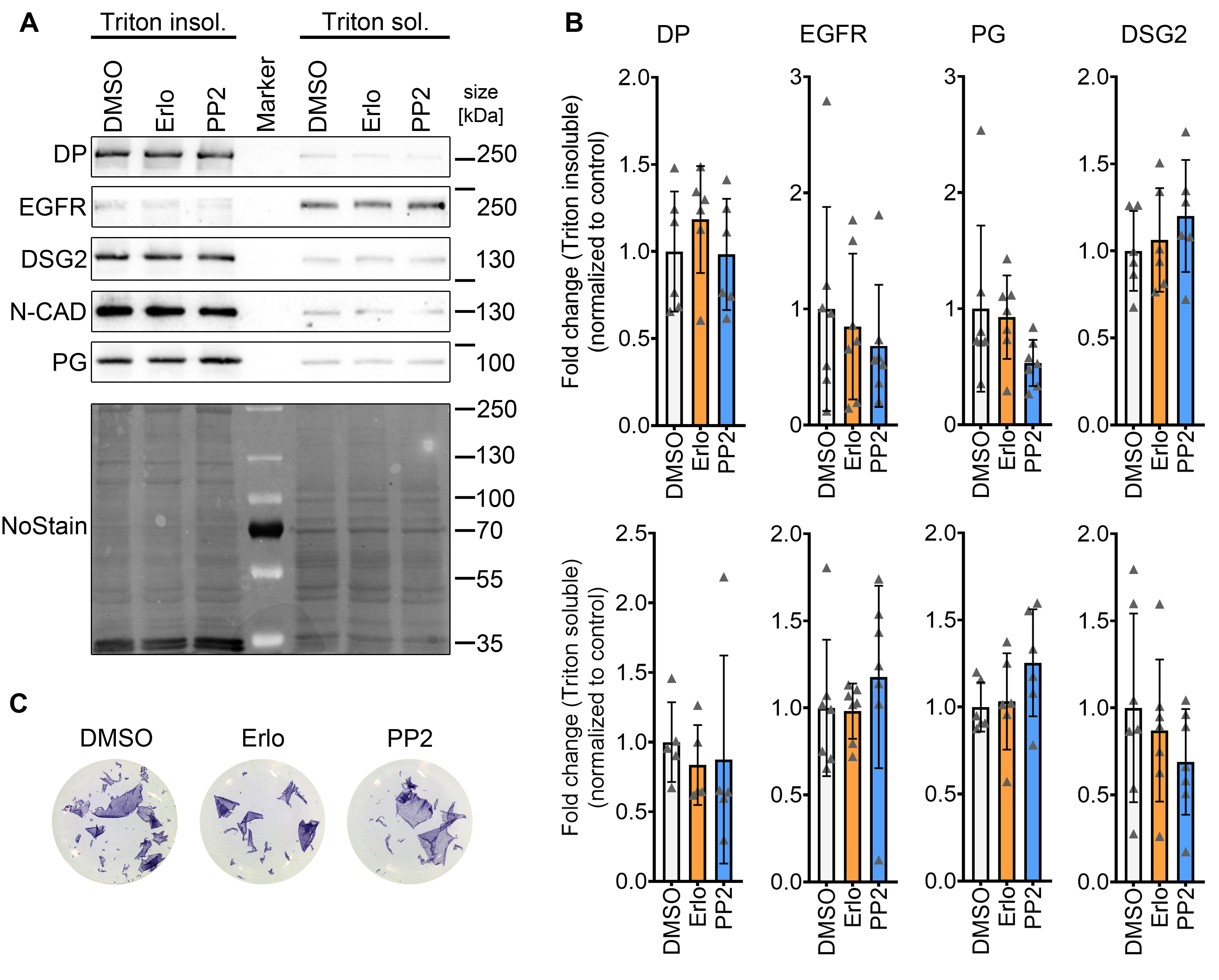

### Supplementary figure 5

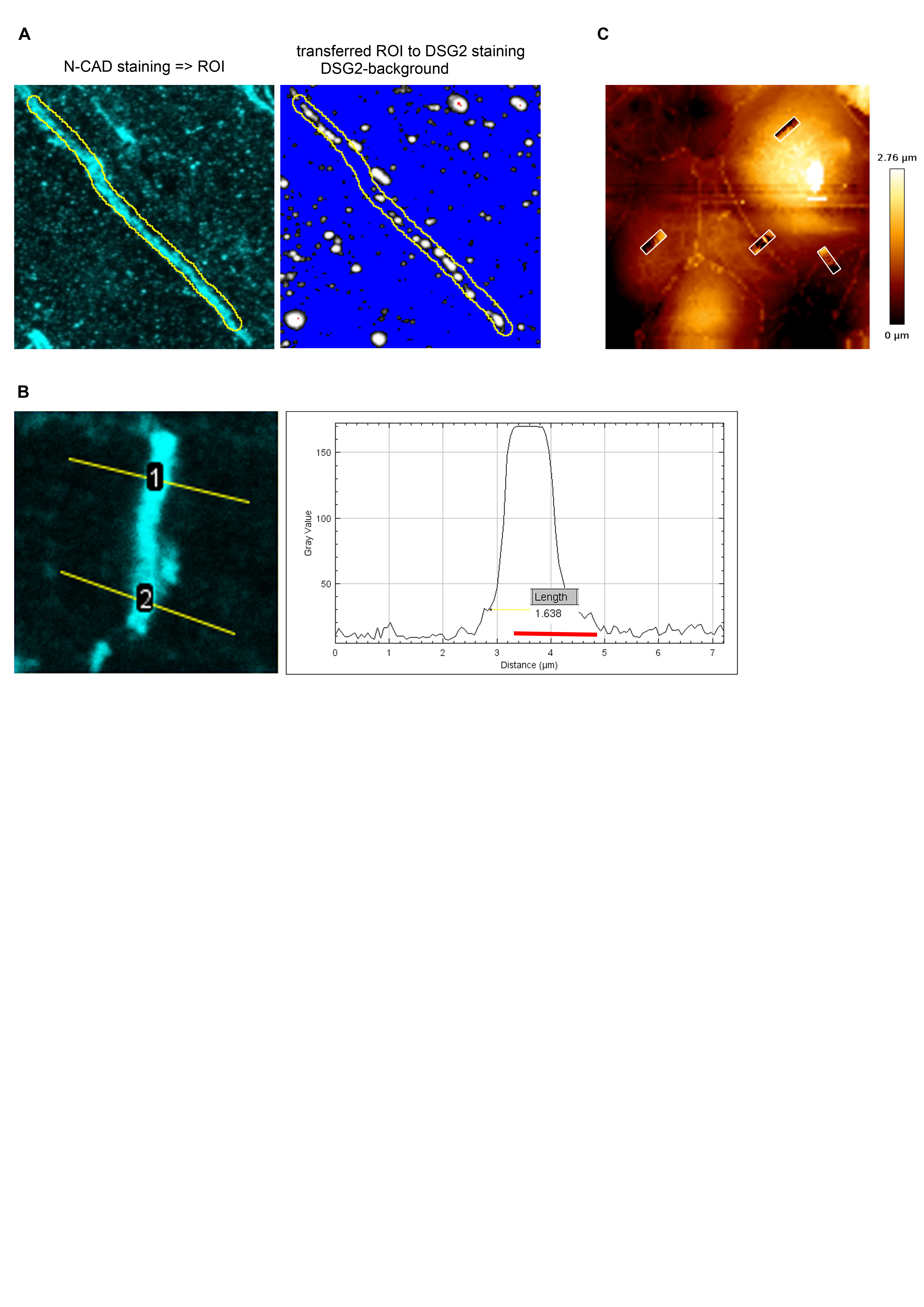
